# An innate granuloma eradicates an environmental pathogen using *Gsdmd* and *Nos2*

**DOI:** 10.1101/2023.03.07.531568

**Authors:** Carissa K. Harvest, Taylor J. Abele, Chen Yu, Cole J. Beatty, Megan E. Amason, Zachary P. Billman, Morgan A. DePrizio, Carolyn A. Lacey, Vivien I. Maltez, Heather N. Larson, Benjamin D. McGlaughon, Daniel R. Saban, Stephanie A. Montgomery, Edward A. Miao

**Affiliations:** Department of Integrative Immunobiology, Duke University School of Medicine, Durham, NC, USA; Department of Molecular Genetics and Microbiology, Duke University School of Medicine, Durham, NC, USA; Department of Microbiology and Immunology, University of North Carolina at Chapel Hill, Chapel Hill, NC, USA; Department of Ophthalmology, Duke University School of Medicine, Durham, NC, USA; Department of Pathology, University of North Carolina at Chapel Hill, Chapel Hill, NC, USA; Department of Cell Biology, Duke University School of Medicine, Durham, NC, USA

## Abstract

Granulomas often form around pathogens that cause chronic infections. Here, we discover a novel granuloma model in mice. *Chromobacterium violaceum* is an environmental bacterium that stimulates granuloma formation that not only successfully walls off but also clears the infection. The infected lesion can arise from a single bacterium that replicates in the presence of a neutrophil swarm. Bacterial replication ceases when macrophages organize around the infection and form a granuloma. This granuloma response is accomplished independently of adaptive immunity that is typically required to organize granulomas. The *C. violaceum*-induced granuloma requires at least two separate defense pathways, gasdermin D and iNOS, to maintain the integrity of the granuloma architecture. These innate granulomas successfully eradicate *C. violaceum* infection. Therefore, this new *C. violaceum*-induced granuloma model demonstrates that innate immune cells successfully organize a granuloma and thereby eradicate infection by an environmental pathogen.

## INTRODUCTION

*Chromobacterium violaceum* is a Gram-negative bacterium found in freshwater sediment and soils. This colorful bacterium, which produces a violet pigment, encodes a type III secretion system (T3SS) that is similar to the invasion-associated T3SS found in *Salmonella* species^1^. *C. violaceum* uses this T3SS to invade and replicate in nonphagocytic cells^2,3^. Immunocompetent people are likely often exposed to *C. violaceum*, but almost never develop symptomatic infection. Conversely, immunocompromised patients can be highly susceptible to *C. violaceum* infection. For example, patients with chronic granulomatous disease, who have defects in the phagocyte NADPH oxidase, are highly susceptible to disseminated *C. violaceum* infection with a mortality rate of around 55%^1,4^. Concomitantly, mice lacking the NADPH oxidase succumb to low dose *C. violaceum* infection within one day^3^. In addition, we previously discovered that resistance to *C. violaceum* requires the T3SS-sensing NLRC4 inflammasome, caspase-1, and gasdermin D^3,5^. Mice deficient in these pyroptosis-inducing genes succumb to very low dose infection^3^. Therefore, *C. violaceum* has potent virulence potential conferred by its T3SS that is fully counteracted by a functional innate immune system.

This defense requires NADPH oxidase and inflammasomes, however, herein we demonstrate a much more complex innate immune defense must be orchestrated in order to eradicate *C. violaceum*. Large distinct lesions form in the liver of wild type C57BL/6 mice during *C. violaceum* infection^5,6^. Nevertheless, these wild type mice never display visible clinical signs and survive the infection. In this manuscript, we discover that these liver lesions are actually organized granulomas that form in order to eradicate this environmental bacterial pathogen.

Granulomas are an organized aggregation of immune cells that surround a persistent stimulus, the origin of which can be either infectious or noninfectious. Granuloma responses are initiated during infection by pathogens across diverse classes of microorganisms, including bacteria, parasites, fungi, and viruses^7-17^. A defining characteristic of a granuloma is the recruitment and organization of inflammatory macrophages into a layer that surrounds the infection^7-11,14,15,18^. Numerous other immune cell types can be present, most prominently T cells that activate macrophages and are normally essential to organize the granuloma architecture^7,8,10,14,18-22^. Granulomas associated with different infections can have diverse histologic architecture^7-17^. For example, some granulomas contain necrotic cores at their centers, whereas others are composed of intact immune cells^8^. Regardless of the specific histologic architecture of the granuloma, all contain a defining organized ring of macrophages.

Granulomas often form around persistent infectious agents, and granulomas often fail to clear such infections^7,10,19,22,23^. These pathogens often survive within the granuloma, leading many to conclude that the granuloma response is a last resort used against an infection that cannot be cleared. Thus, granulomas are often considered as a way to restrain an infectious agent and prevent its dissemination, albeit while simultaneously being unable to eradicate the infectious agent. Actually, individual granulomas can be heterogeneous – some granulomas clear the infection whereas other granulomas have progressive infection containing viable organisms even within the same animal^24^. Given the diverse clinical outcomes and the lack of understanding as to what drives the formation of various granulomatous responses, novel models are needed to study the range of the granuloma responses, particularly mouse models that leverage advanced immunological technologies^25,26^.

Other granuloma-inducing pathogens, such as *Mycobacterium tuberculosis*, do not clearly demonstrate a critical role for the pyroptotic pathway proteins (inflammasomes, caspase-1, and gasdermin D). Some studies show that murine *M. tuberculosis* infection is exacerbated in the absence pyroptotic pathway mutants^27^, whereas others show no role^28,29^. These contrast with the macaque *M. tuberculosis* infection model, where caspase-1 played a minimal role unless macaques were treated with anti-PD1, when caspase-1 activation was correlated with higher M. tuberculosis burdens^30^. This detrimental role was recapitulated in knockout mice when PD-1 knockouts were crossed with *Casp1*^*–/–*^*Casp11*^*–/–*^ double knockouts^30^. *M. tuberculosis* might be detected by inflammasomes, but also has strategies to evade detection^31-40^. Varying penetrance of evasion in different models or strains could explain these disparate results^41^. Thus, whether pyroptotic proteins are useful in the granuloma response remains unclear.

Herein, we examine the granuloma response to *C. violaceum* from the initiating events through to clearance of the bacterium and provide definitive evidence for a beneficial role of pyroptotic proteins during the granuloma response.

## RESULTS

### *C. violaceum* triggers formation of a necrotic granuloma

Intraperitoneal *C. violaceum* infection of wild type C57BL/6J mice caused macroscopic white-to-cream-colored lesions specifically in the liver (Figure 1A)^5,6^. Lesions were evenly distributed and typically 1-2 mm in diameter at 5 days post infection (dpi) (Figure 1B). We investigated the histologic morphology by hematoxylin and eosin (H&E) staining and consistently observed three distinct layers in every lesion (Figure 1C; green box and 1D). First the center was characterized by dark hematoxylin (purple) staining of amorphous material locking defined cell borders and features, consistent with necrotic debris (Figure 1C).

**Figure 1.**
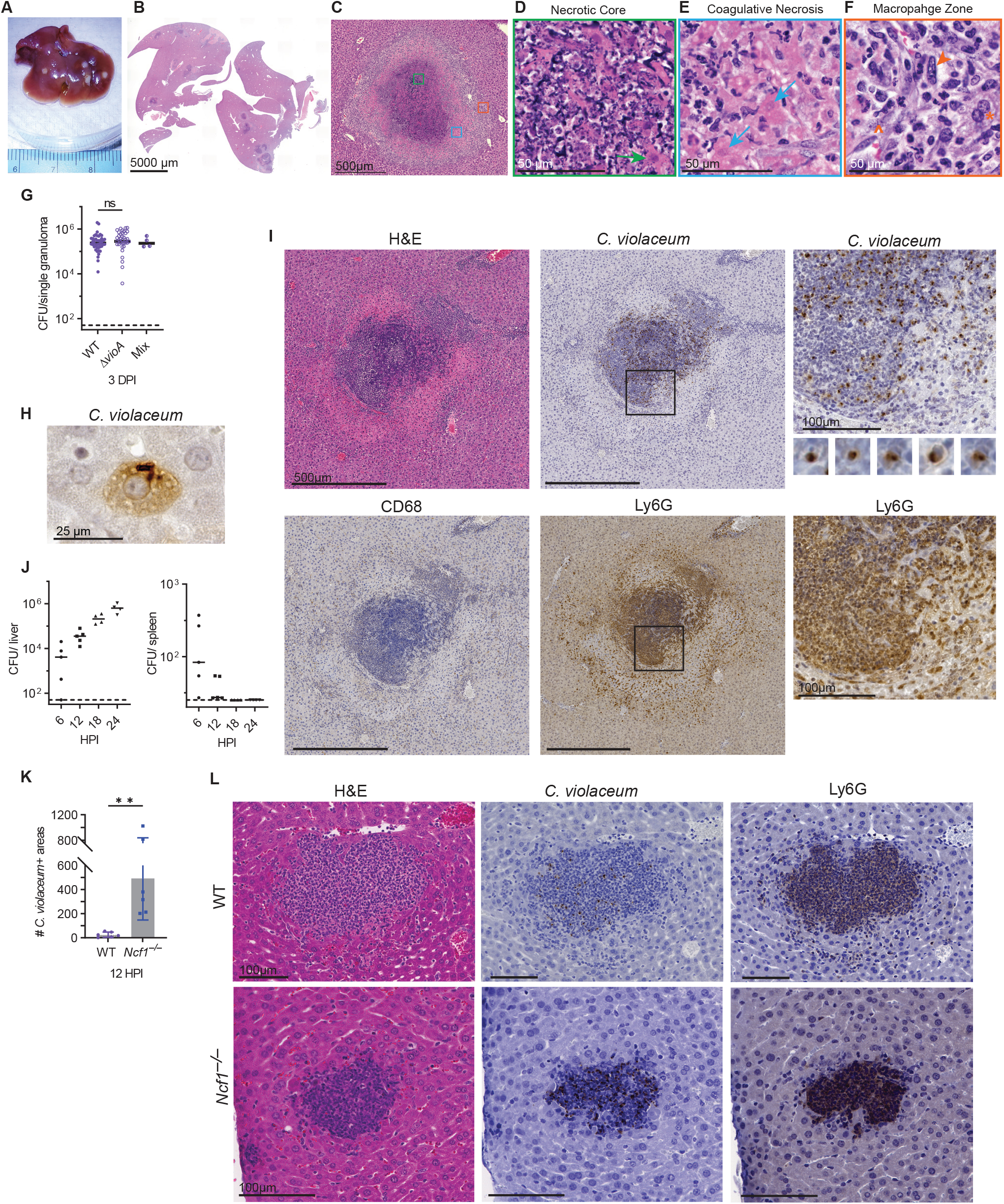
*C. violaceum* replicates despite a neutrophil swarm. (A-L) Mice were infected with WT *C. violaceum* or indicated strains. (A-F) Gross pathology and H&E staining of WT livers 5 dpi. Representative of 10 experiments, each with 3-4 mice, each with multiple granulomas per section. Arrow, coagulative necrotic hepatocyte; arrowhead, macrophage; carrot, fibroblast. (G) Inoculum was a 1:1 mixture of WT and *∆vioA C. violaceum*. 83 individual granulomas were dissected 3 dpi, bacterial burdens determined, and scored for color. Data combined from two experiments. n.s., not significant by Kruskal-Wallis test. (H) Visualization of *C. violaceum* within a hepatocyte at 12 hpi by IHC. (I) Serial sections stained by H&E or indicated IHC markers 1 dpi. Representative of 5 experiments, each with 4 mice, each with multiple granulomas per section. (J) Bacterial burdens in the liver and spleen at indicated timepoints. Representative of 2 experiments; each point is a single mouse. (K) Quantitation of *C. violaceum*-positive stained areas per section 12 hpi in WT and *Ncf1*^*–/–*^ mice. Combined data from 2 experiments, each with 3 mice per genotype. **p<0.01, by unpaired two-tailed t test (L) Serial sections stained by H&E or indicated IHC markers 12 dpi in WT and *Ncf1*^*–/–*^ mice. Representative of 2 experiments each with 3-4 mice per genotype. Dashed line, limit of detection; solid line, median.

Surrounding this necrotic core was the second layer, composed of degenerative cells that retained cell borders but lost morphologic subcellular detail, identified as hepatocytes based on their location, size, and nuclear morphology. (Figure 1C and 1E). The nuclei retained their size and shape but displayed faded hematoxylin chromatin staining, and with some having an absent nucleus (degeneration). The cytoplasm stained brightly eosinophilic (pink), hepatocyte boarders remained apparent, and the cells sporadically merged with the necrotic core (Figure 1D and 1E), indicative of coagulative necrosis^42^.

The third layer, exterior to the layer of coagulative necrosis, was predominantly composed of cells with abundant cytoplasm and a large, oval nucleus with open chromatin which is consistent with activated macrophages (Figure 1F). We refer to this layer as the “macrophage zone”. Within this macrophage zone, scattered fibroblasts were present, identified by their fusiform shape, faint eosinophilic cytoplasmic staining, and a distinct, dark elongated nucleus (Figure 1F; carrot). Immediately exterior to the macrophage zone are viable hepatocytes at 5 dpi (Figure 1F; asterisk). The distinct, organized zone of macrophages surrounding each lesion is a defining feature of a granuloma, with the presence of a necrotic core subcategorizing them as a necrotic granuloma.

### A granuloma forms in response to one bacterium

To determine whether each granuloma was initiated by a single bacterium, we infected mice with a 1:1 ratio of WT (violacein positive) and *∆vioA* mutant (violacein negative) *C. violaceum* strains and determined the number and color of colony forming units (CFUs) from dissected single granulomas (Figure 1G and S1A). The vast majority of single granulomas contained only one *C. violaceum* strain (Figure 1G and S1A). The mixed color granulomas may arise from errors in dissection where two granulomas where harvested instead of one. This result suggests that these granulomas are seeded by a single bacterium. In addition, WT and *∆vioA* seeded equivalent numbers of granulomas with similar burdens per granuloma (Figure 1G), indicating that violacein pigment is dispensable for liver infection^43^. This suggests that a single bacterium replicates to a median 5.4×10^6^ CFUs in each granuloma by 3 dpi. Clearly, the innate immune response fails to halt *C. violaceum* replication within the first 3 dpi.

### Neutrophil swarming fails to eradicate *C. violaceum*

We next investigated the early hours of infection to understand the granuloma development. *C. violaceum* infects hepatocytes using its T3SS^2^, a cell type in which we were unable to detect caspase-1 protein^3^, suggesting that this niche is not defended by caspase-1. We visualized *C. violaceum* by immunohistochemical staining 12 hours post infection (hpi) and observed bacteria within hepatocytes (Figure 1H). Other bacteria were also observed in immune cells (Figure S1B). We observed multiple bacteria within each hepatocyte, suggesting that the bacteria were replicating by 12 hpi. At 24 hpi, larger, distinct lesions appear, comprised of a swarm of neutrophils identified morphologically by their multilobulated nucleus, and confirmed by Ly6G staining (Figure 1I). Macrophages were absent from these lesions by morphological identification and verified by CD68 staining (Figure 1I and S1C). *C. violaceum* staining showed bacteria only within the neutrophil swarm and absent from surrounding hepatocytes (Figure 1I). Bacteria were tightly clustered within individual neutrophils (Figure 1I). Bacterial burdens in the liver increased by ∼100-fold between 6 hpi and 24 hpi (Figure 1J). In contrast, the spleen had low burdens at 6 hpi, and burdens were cleared by 18 hpi (Figure 1J) and remain sterile at 3 dpi^3^. Thus, despite the presence of a neutrophilic response within the first 24 hpi, *C. violaceum* continues to replicate exponentially within the liver.

Although the histologically apparent neutrophils failed to clear the bacteria, reactive oxygen species (ROS) are required to slow bacterial replication over the first 24 hpi. Myeloid cells generate ROS in the phagosome using the phagocyte NADPH oxidase, which includes p47^*phox*^ encoded by *Ncf1. Ncf1*^*–/–*^ mice succumb to even 100 CFUs of *C. violaceum* within 24 hpi^3^. *Ncf1*^*–/–*^ mice showed increased numbers of *C. violaceum*-positive liver lesions at 12 hpi (Figure 1K). These ranged from a few neutrophils to sizable neutrophil swarms, albeit smaller than those seen in WT mice (Figure 1L). Again, *C. violaceum* staining clustered tightly within the neutrophil swarm in *Ncf1*^*–/–*^ mice (Figure 1L). Thus, although the NADPH oxidase limits seeding of lesions by *C. violaceum*, neutrophils nevertheless fail to eradicate *C. violaceum* which continues to grow within the neutrophil swarm of WT mice.

### A granuloma forms around the infected neutrophil swarm

Because neutrophils fail to eradicate *C. violaceum*, a more effective immune response is needed. By 3 dpi, the lesions have enlarged (Figure 2A). The neutrophils observed at 1 dpi now form the bulk of the necrotic core at 3 dpi, identified by Ly6G staining (Figure 2B and 2C). We observed a zone of coagulative necrotic hepatocytes surrounding the necrotic core (Figure 2B). *C. violaceum* staining was observed in the necrotic core, but not inside the sparse coagulative necrotic hepatocytes in the core (Figure 2D; carrots). At 3 dpi we observed a thin macrophage zone identified by morphology and confirmed with CD68 staining (Figure 2B and 2C). Bacterial staining was essentially absent in the macrophage zone, except rare areas where neutrophils were infiltrating the periphery of the granuloma (Figure 2B and 2C). Further, no *C. violaceum* staining was observed in the healthy hepatocytes outside the granuloma (Figure 2B and 2C). This architecture indicates the formation of an early granuloma which can be subcategorized into a necrotic suppurative granuloma (pyogranuloma)^8^. At 5 dpi, the macrophage zone became thicker, with pronounced CD68 staining (Figure 2B and 2C), while neutrophil staining at the periphery became sparser (Figure 2B and 2C).

**Figure 2.**
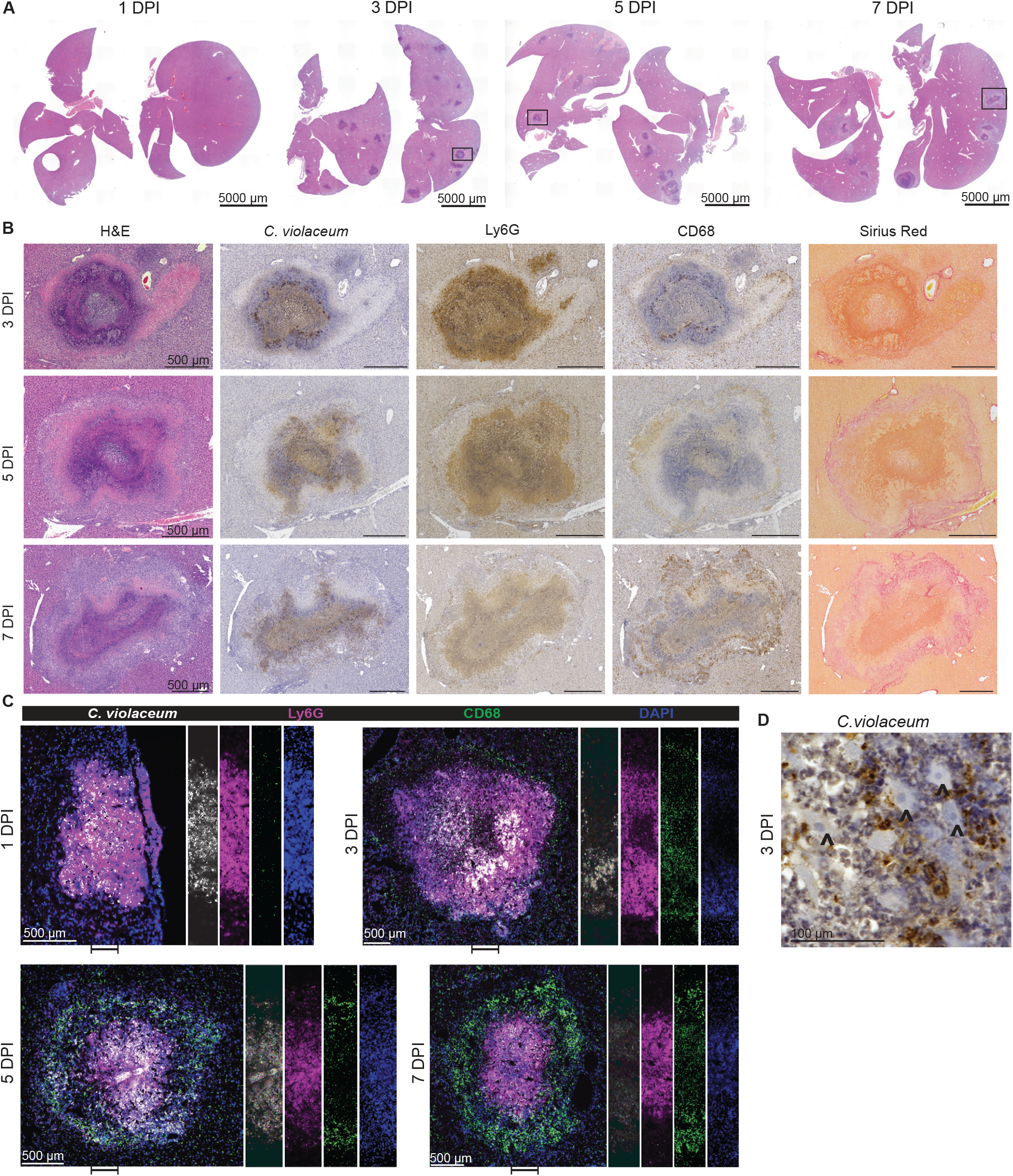
A granuloma forms around the infected neutrophil swarm. (A-D) Mice were infected with WT *C. violaceum*. (A) H&E staining of WT livers 1, 3, 5, and 7 dpi. Representative of 10 experiments (3 and 5 dpi) and 5 experiments (1 and 7 dpi), each with 3-4 mice, each with multiple granulomas per section. (B) Serial sections stained by H&E or indicated IHC markers 3, 5, and 7 dpi. Representative of 10 experiments (3 and 5 dpi) and 5 experiments (7 dpi), each with 4 mice, each with multiple granulomas per section. (C) WT liver sections stained with indicated IF markers 3, 5, and 7 dpi. Representative of 2 experiments; each with 4 mice, each with multiple granulomas per section. (D) Visualization of *C. violaceum* 3 dpi; carrot, coagulative necrotic hepatocytes.

Concomitantly, we identified fibroblasts by morphology and colocalized collagen deposition, both within the macrophage zone (Figure 2B and 2C). *C. violaceum* staining was still predominantly within the necrotic core (overlapping with Ly6G); notably, all the bacterial staining was contained within the granuloma (Figure 2B and 2C). At 7 dpi the macrophage zone had become even thicker while the necrotic core appears smaller than at 3 and 5 dpi (Figure 2A, 2B, and 2C). At this timepoint the coagulative necrosis zone persists, and collagen staining is also more prominent (Figure 2B). Epithelioid macrophages are present in the granulomas of other animal models and express E-cadherin to form tight junctions, functioning to help wall off the pathogen ^44^. However, we did not observe E-cadherin staining in the *C. violaceum*-induced granuloma (Figure S1D). This *C. violaceum*-induced granuloma model demonstrates the ability of macrophages to organize a granuloma that successfully prevents bacterial dissemination.

At 14 dpi most granulomas were visibly smaller, although rare granulomas remain large in occasional mice (Figure 3A). In a typical 14 dpi granuloma the macrophage zone remained prominent, while the necrotic core became even smaller; in some granulomas the necrotic core was completely absent (Figure 3B and S1E; arrows). The coagulative necrotic zone remained prominent, although the cellular borders of the coagulative necrotic hepatocytes were no longer distinct. The macrophage zone remained prominent and began to infiltrate the coagulative necrotic zone (Figure 3B). At earlier timepoints, *C. violaceum* staining and CD68 staining do not overlap (Figure 2B and 2C), however, surprisingly at 14 dpi these stains now overlap in the macrophage zone (Figure 3B and S1F). This staining pattern suggests that dead bacterial antigens have been phagocytosed by macrophages in the process of removing the necrotic core. In resolved livers, which were common at 21 dpi, we observed small clusters of various immune cells and small, flat areas of collagen (Figure 3C and 3D). Such areas are consistent with a granuloma that has contracted and has almost completely resolved.

**Figure 3.**
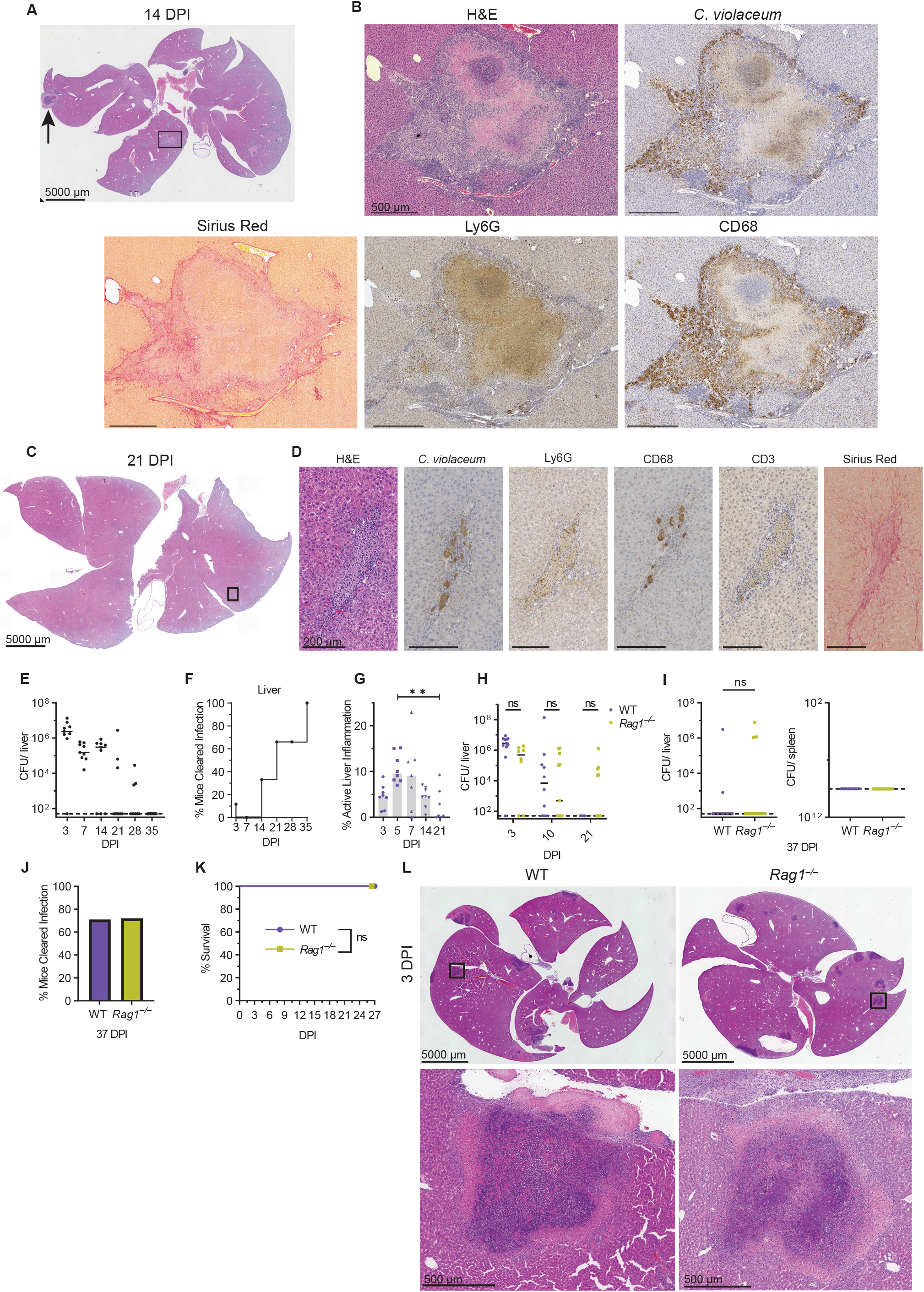
Adaptive immunity is not required to resolve the granuloma. (A-L) Mice were infected with WT *C. violaceum*. (A) H&E staining of WT livers 14 dpi. Representative of 3 experiments, each with 3-4 mice, each with multiple granulomas per section. (B) Serial sections stained by H&E or indicated IHC markers 14 dpi. Representative of 3 experiments, each 3-4 mice, each with multiple granulomas per section. (C) H&E staining of WT livers 21 dpi. Representative of 3 experiments, each with 3-4 mice, each with multiple granulomas per section. (D) Serial sections stained by H&E or indicated IHC markers 21 dpi. Representative of 3 experiments, each 3-4 mice, each with multiple granulomas per section. (E, H, and I) Bacterial burdens in the liver and spleen at indicated timepoints. Representative of 2 experiments; each point is a single mouse. ns, not significant, by Two-way ANOVA (H) or Kruskal-Wallis test (I). (F) Bacterial burdens from (E) displayed as % bacterial clearance. (G) Percent active inflammation quantified per whole liver section stained with H&E. Data combined from 2 experiments, each point is a single mouse. **p<0.01, by Kruskal-Wallis test. (J) Bacterial burdens from (I; liver) displayed as % bacterial clearance. (K) Survival analysis of WT and *Rag1*^*–/–*^ mice. Representative of 2 experiments each with 8-12 mice per genotype. ns, not significant, by Kaplan-Meier survival analysis. (L) H&E staining of WT and *Rag1*^*–/–*^ livers at 3 dpi. Representative of 2 experiments, each with 3-4 mice, each with multiple granulomas per section. Dashed line, limit of detection; solid line, median.

### Individual granulomas resolve at different rates

Maximal bacterial burdens peak at 3 dpi and decrease over time thereafter (Figure 3E and S2A). This correlates with the appearance of the macrophage zone, suggesting that the switch of neutrophils to macrophages halts bacterial replication. Starting at 7 dpi, some mice cleared the infection, and the proportion of mice that cleared steadily increased over time, although the kinetics of clearance vary slightly between experiments (Figure 3E, 3F, S2A and S2B). The majority of mice clear the infection between 14 and 35 dpi (Figure 3F and S2A).

Infrequently at later time points, sporadic mice still had bacterial burdens. We hypothesize these burdens arise from single granulomas that have not yet resolved (Figure 3A). In such remaining granulomas, *C. violaceum* staining is still located in the necrotic core (Figure S2C), more similar to a typical granuloma at 5 dpi. The amount of total liver inflammation decreases over time with visual disappearance of the necrotic core (Figure 3G and S2E). This suggests that individual granulomas eliminate bacteria independently of each other within the same mouse. Indeed, when we dissected single granulomas at 7 dpi some granulomas were sterile while others contained burdens within the same mouse, and a greater proportion became sterile at 10 dpi (Figure S2E). These observations suggest that the granuloma response successfully clears *C. violaceum* infection.

### Adaptive immunity is not required

Most granuloma responses involve the adaptive immune system in which T cells are maintain or support the organization of the macrophage zone^7,8,14,21,22^. However, in the *C. violaceum*-induced granuloma T cells within the macrophage zone were sporadic and unorganized (Figure 3D and S2F). Stochastic clearance of burdens began at 10 dpi in both WT and *Rag1*^−/−^ mice (Figure 3E, 3F, 3G, and 3H). The proportion of mice that cleared the infection increased through 37 dpi when both WT and *Rag1*^−/−^ had the same percentage of mice sterilizing the infection (Figure 3I and 3J). The trend of clearance between WT and *Rag1*^−/−^ mice is similar, which is consistent with previous results using a lower dose infection^3^. Furthermore, both groups survived the infection equally (Figure 3K). The abundance and the architecture of granulomas per liver section between WT and *Rag1*^−/−^ was similar at 3 dpi (Figure 3L) and of 3 *Rag1*^−/−^ mice analyzed by H&E staining at 26 dpi, two had no granulomas and one had a single small resolving granuloma (Figure S2G). Although adaptive immunity is not required for this granuloma response, this does not preclude adaptive responses from being critical in future infections. Indeed, WT mice that were rechallenged with *C. violaceum* did not develop visible granulomas or bacterial burdens at 3 days post-secondary infection (not shown). Taken together, *C. violaceum* induces the formation of innate granulomas that organize, eradicate the bacteria, and resolve without the need for adaptive immunity.

### Ischemia and coagulation outside the granuloma

We observed other pathology outside the granuloma structure. Ischemia was observed in WT mice at 3 and 5 dpi (Figure 2A, arrowhead, and S3A) often directly adjacent to a granuloma (Figure S3B and S3C), suggesting that the granuloma occluded or affected the blood supply. Ischemia observed in isolation could be adjacent to a granuloma outside the histologic plane. Ischemic areas were of various sizes and did not stain for *C. violaceum* antigens (Figure S3D). We did not observe ischemia past 7 dpi suggesting rapid regeneration, consistent with the robust regenerative power of hepatocytes. We frequently observed thrombosis in the liver vasculature, some of which occluded hepatic vessels and caused ischemia in the downstream tissue (Figure S3C and S3E).

### Perforin and caspase-7 are not required for the granuloma response

Natural killer (NK) cell attack reduces *C. violaceum* burdens in the liver^3^ requiring perforin and caspase-7^5^. We have shown cleaved caspase-7 staining in what we now know are granulomas of WT mice^5^, however, NK cells were scarce and unorganized within the macrophage zone (Figure S3F). *Prf1*^*–/–*^ mice survive and clear *C. violaceum* infection by 19 dpi (Figure S3G and S3H), and both *Prf1*^*–/–*^ and *Casp7*^*–/–*^ mice have similar granuloma architecture compared to WT mice (Figure S3I). Thus, although some NK cells are present, neither perforin nor caspase-7 are required for the overall granuloma-mediated clearance of *C. violaceum*.

### Gasdermin D is essential during the granuloma response

*C. violaceum* uses a T3SS to reprogram host cells. However, the activity of this T3SS can be detected by the NAIP/NLRC4 inflammasome, activating caspase-1, which cleaves pro-IL-1β, pro-IL-18, and gasdermin D to their active forms. Once cleaved, gasdermin D oligomerizes and forms a pore that inserts into the inner leaflet of the plasma membrane, resulting in the influx of fluids causing the cell to swell and undergo pyroptotic lysis. Pyroptosis is an inflammatory form of regulated cell death that can combat intracellular infection^45^.

*Casp1/11*^−/−^ and *Gsdmd*^*–/–*^ mice infected with *C. violaceum* succumb to infection 3-9 dpi and had increased burdens in both the liver and spleen (Figure 4A, 4B, and 4C)^3,5^. We observed obvious abnormalities within the granuloma architecture of both knockouts at 3 dpi (Figure 4D and 4E), with complete loss of the distinct layers seen in WT mice (Figure 4F). Most lesions in these knock out mice have increased abundance of necrosis containing Ly6G and *C. violaceum* staining (Figure 4F and 4G). These lesions have a ‘budding’ morphology with strong Ly6G staining but little CD68 staining (Figure 4F and 4G; H&E, brackets). Within each budding area and throughout each lesion in *Casp1/11*^−/−^ and *Gsdmd*^*–/–*^ mice we observed *C. violaceum* staining that extended to the outer edge of the lesion (Figure 4F and 4G), and outside the defective granulomas (Figure S3J and S3K). This loss of bacterial containment could arise by escape from the defective granuloma or seeding via portal circulation from the infected spleen. These data clearly demonstrate that caspase-1/11 and gasdermin D are required for the granuloma to form and contain *C. violaceum* infection.

**Figure 4.**
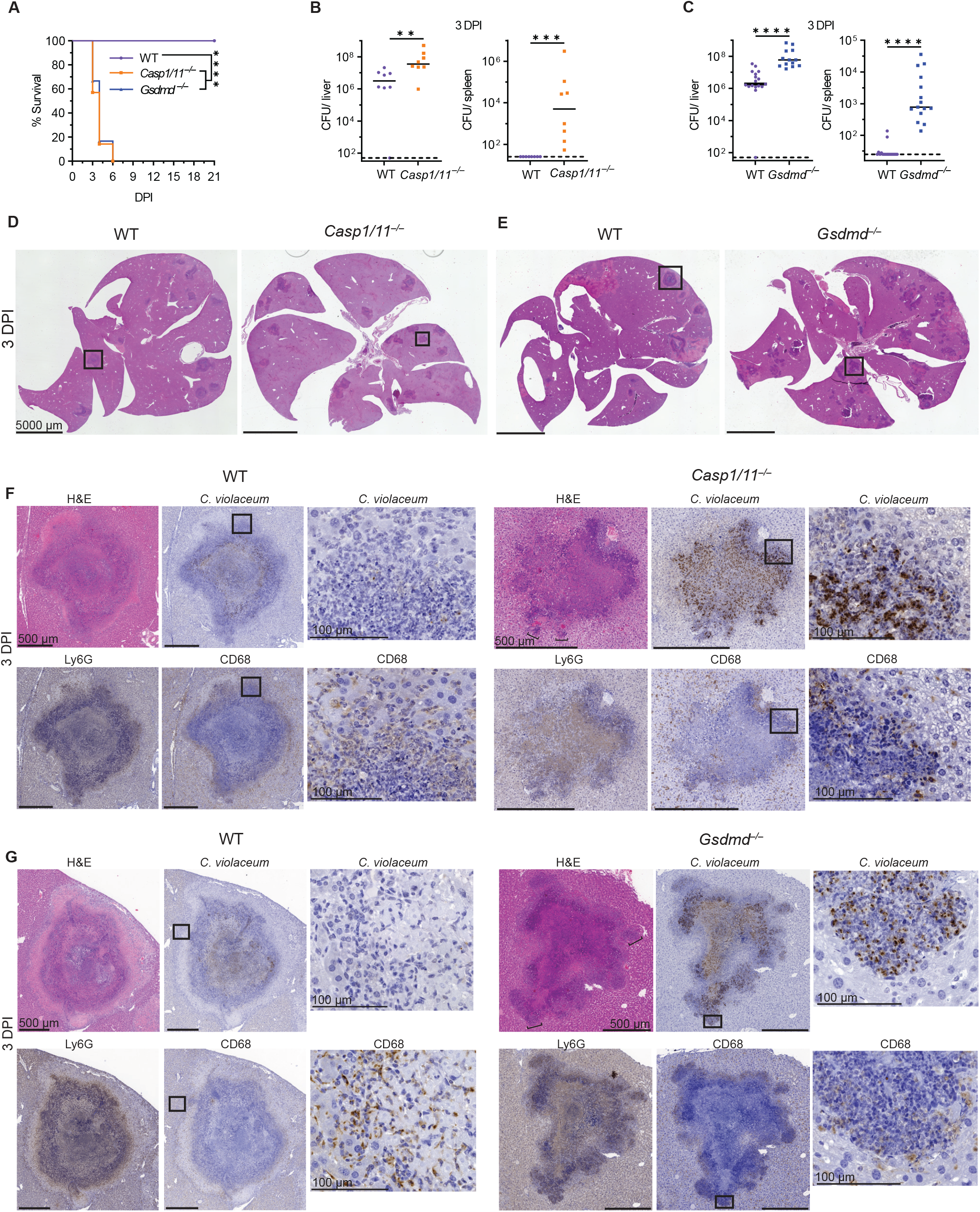
Gasdermin D is essential during the granuloma response. (A-G) Mice were infected with WT *C. violaceum*. (A) Survival analysis of WT, *Casp1*^*–/–*^*Casp11*^*–/–*^ (*Casp1/11*^*–/–*^), and *Gsdmd*^*–/–*^ mice. Representative of 2 experiments, each with 6-8 mice per genotype. ****p<.0001, by Kaplan-Meier survival analysis with Bonferroni correction for multiple comparisons. (B and C) Bacterial burdens in the liver and spleen at 3 dpi. Data combined from 2 experiments, each point is a single mouse. **p<0.01, ***p<0.001, ****p<.0001, by Mann-Whitney test. (D-G) Serial sections stained by H&E or indicated IHC markers 3 dpi. Representative of 2 experiments, each 3-4 mice, each with multiple granulomas per section. Dashed line, limit of detection; solid line, median.

### Spatial mapping of the granuloma transcriptome

To investigate gene expression within different zones of the granuloma, we used spatial transcriptomics (10x Genomics Visium) to capture mRNA from granulomas in WT mice during *C. violaceum* infection (Figure S4A). Liver sections at 12 hpi, 1, 3, 5, 10, 14 and 21 dpi contained granulomas that appear representative of each timepoint. Although the 7 dpi liver section unfortunately contained a non-representative granuloma that was smaller than normal (Figure S4A), this timepoint was nevertheless included within the analysis. We identified sixteen distinct expression clusters within the granuloma and the surrounding liver (Figure S4A). Clusters had distinct distribution profiles within the layers of the granuloma architecture (Figure 5A and S4B). To assign cell types within each capture area we used published single cell sequencing data from the Mouse Cell Atlas (Figure 5A, 5B, S4A, and S4B).

**Figure 5.**
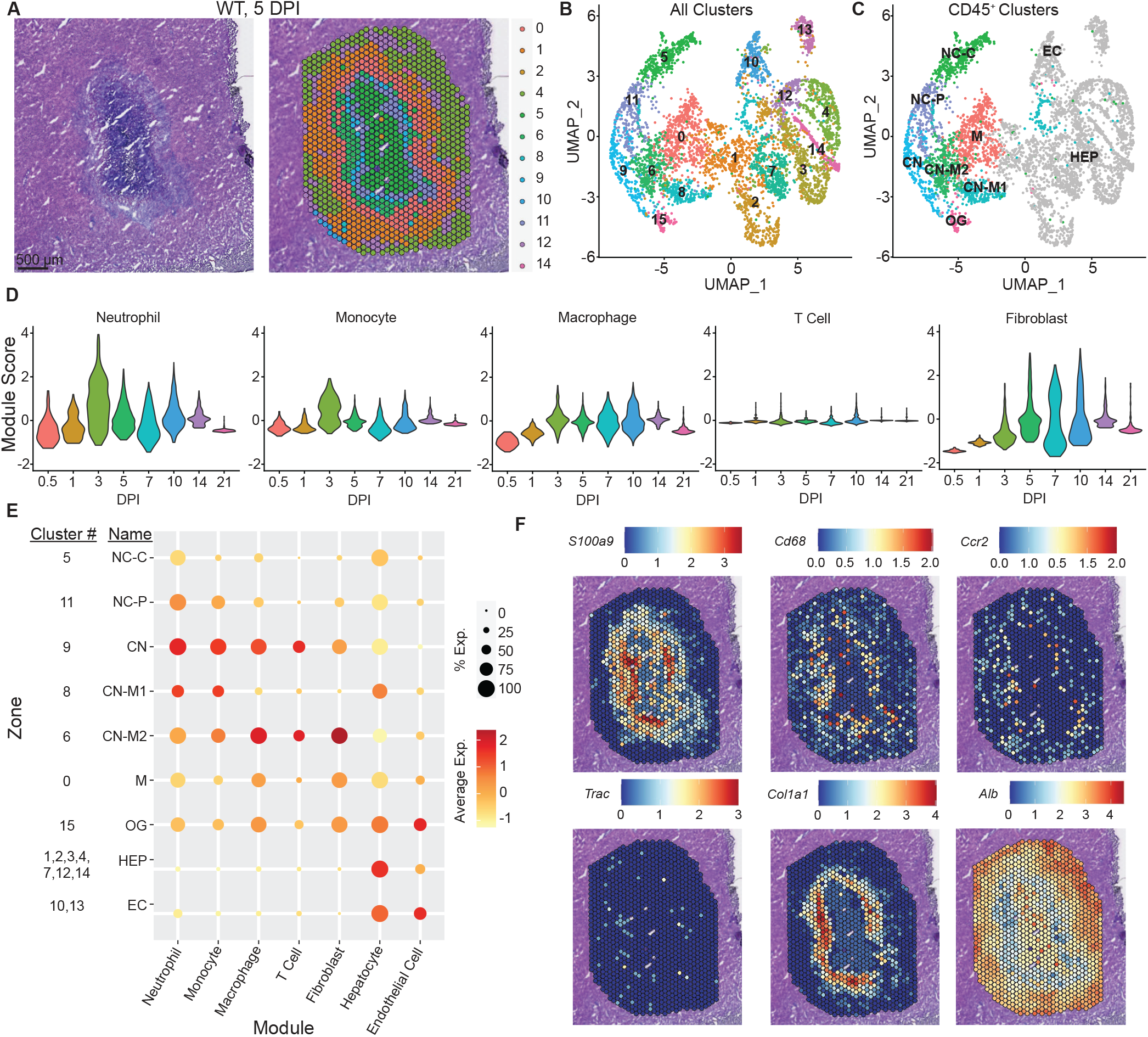
Spatial mapping of the granuloma transcriptome. (A-F) Mice were infected with WT *C. violaceum* and one liver was harvested per indicated timepoint for spatial transcriptomic analysis. (A) H&E and spatial transcriptomic orientation of indicated clusters from same granuloma at 5 dpi. (B) UMAP plot for cluster expression orientation for all clusters from all timepoints of infection. (C) UMAP plot of cluster expression in designated locations within granuloma architecture. (D) Module scores of various immune cell and fibroblast expression markers from granulomas throughout infection time course. (E) Indicated cell type expression dot plot in each cluster location within the granuloma architecture of all combined time points of infection. (F) Spatial transcriptomic expression of indicated genes representing immune and non-immune cells within 5 dpi granuloma. *S100a9* (neutrophils), *Cd68* (macrophages), *Ccr2* (monocytes), *Trac* (T cells), *Col1a1* (fibroblasts), and *Alb* (hepatocytes and endothelial cells).

Immune cell-enriched *Ptprc* (CD45)-positive areas were located on the left half of the UMAP plot, while non-immune clusters were on the right and generally expressed the hepatocyte marker *Alb* (Figure S4C and S4D). We annotated clusters by both sequencing- and histological-based parameters: necrotic core (NC) – identified based on their basophilic properties and low RNA abundance; coagulative necrotic zone (CN) – identified based on spatial overlap with regions of diffuse tissue damage surrounding the necrotic core; CN-macrophage zone (CN-M) – identified based on their presence at the nexus between the CN and M zones; macrophage zone (M) – identified based on the mildly basophilic nature of the underlying tissue and presence outside the CN zone (Figure 5C and S4E). Sub-zonal annotation was used to denote unique clusters found in the same zone. We created gene expression modules using markers from published single cell RNA sequencing datasets from mouse liver^46,47^ cross referenced in ImmGen for cell-type-specific expression, and selected marker genes expressed in our spatial transcriptomics dataset. We created modules for neutrophils, monocytes, macrophages, T cells, fibroblasts, hepatocytes, and endothelial cells (Figure S4G). The neutrophil module was peaking at 3 dpi, after which their expression diminishes but remains present over time (Figure 5D). Monocytes prevalence peaked at 3 dpi and declined thereafter (Figure 5D), suggesting that the monocytes differentiated into macrophages, which became maximal at 3 dpi and stayed steady throughout the granuloma response (Figure 5D). Consistent with their lack of involvement in the granuloma response, T cell gene expression was minimal (Figure 5D). Fibroblast gene expression peaks at 5 dpi and remains steady until later timepoints (Figure 5D).

By assigning modules to zones we observed that neutrophil gene expression was present primarily within the necrotic core and the coagluative necrotic layers (Figure 5E), which histologically are composed of necrotic debris, indicating that mRNAs from dead neutrophils persists. Macrophage and monocyte gene expressions were present primarily throughout the coagulative necrotic zone, macrophage zone, and just outside the macrophage zone (Figure 5E) but were minimal in the necrotic core. Fibroblast gene expression was seen throughout the macrophage zone spreading into the coagulative necrotic zone (Figure 5E), consistent with Sirius red staining (Figure 2B). Hepatocyte and endothelial cell gene expression was diminished within the granuloma and was predominant in the outer surrounding tissue (Figure 5E). Using key genes from each cell module, we confirmed their location within the granuloma by spatial expression in each capture area (Figure 5F). Collectively, this spatial transcriptomics data confirms the histologically visualized distribution of cell types and the specific staining for cell types with marker-specific antibodies.

### Nitric oxide is required to clear *C. violaceum* in the granuloma

*Nos2* and *Acod1* both have been previously implicated in other granuloma models^48-52^ and we saw expression of both in our spatial transcriptomics data. *Nos2* encodes inducible nitric oxide synthase (iNOS), an enzyme that produces nitric oxide (NO) that can kill bacteria. *Acod1* encodes IRG1, an enzyme that produces itaconate, which can also kill bacteria and immunometabolic activities^53,54^. Indeed, we observed that NO can kill *C. violaceum* in vitro, and although itaconate alone was not toxic, it enhanced NO toxicity (Figure S5A). This suggested that iNOS and IRG1 might cooperate to sterilize the granuloma. However, *Acod1*^*–/–*^ mice survived *C. violaceum* infection with normal bacterial burdens (Figure S5B and S5C). Therefore, IRG1 is expressed in the granuloma, but is not essential for clearance of *C. violaceum*.

In contrast, all *Nos2*^*–/–*^ mice succumbed to *C. violaceum* infection (Figure 6A), confirmed with littermate controls (Figure S5D). The importance of iNOS was not yet manifested at 3 dpi, when *C. violaceum* continued to replicate normally in the liver and continued to be cleared from the spleen of *Nos2*^−/−^ mice (Figure 6B), suggesting that iNOS did not act during the neutrophil swarm phase of the infection. Bacterial burdens increased in the liver of *Nos2*^*–/–*^ mice at 5 dpi and continued to increase at 7 dpi compared to WT mice, and *Nos2*^−/−^ mice succumbed shortly thereafter (Figure 6A, 6C, and 6D). The spleen also showed high *C. violaceum* burdens for the majority of the *Nos2*^*–/–*^ mice at these later time points (Figure 6C and 6D), which may reflect dissemination from the liver rather than an open niche in the spleen.

**Figure 6.**
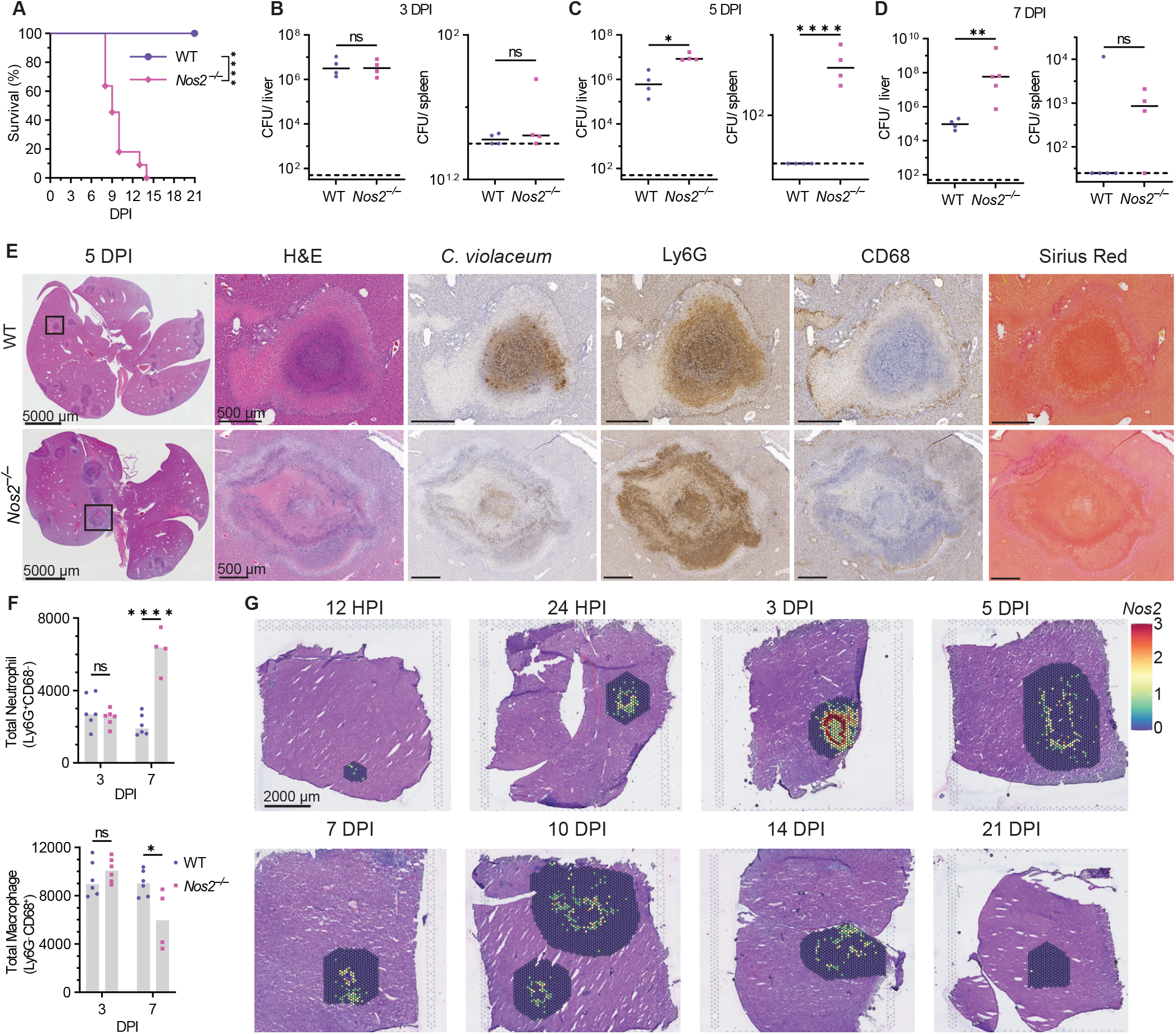
Nitric oxide is required to clear *C. violaceum* in the granuloma. (A-G) Mice were infected with WT *C. violaceum*. (A) Survival analysis of WT and *Nos2*^*–/–*^ mice. Representative of 2 experiments, each with 11 mice per genotype. ****p<.0001, by Kaplan-Meier survival analysis. (B-D) Bacterial burdens in the liver and spleen at 3, 5, and 7 dpi. Data representative of 2 experiments, each point is a single mouse. Dashed line, limit of detection; solid line data median. ns, not significant, *p<0.05, **p<0.01, ****p<.0001, by Unpaired two tailed t test (liver B; spleen C; liver and spleen D) and Mann-Whitney test (spleen B; liver C). (E) Serial sections stained by H&E or indicated IHC markers of WT and *Nos2*^*–/–*^ livers and granulomas 5 dpi. Representative of 2 experiments, each with 3-4 mice, each with multiple granulomas per section. (F) Flow cytometry of total neutrophil and macrophage numbers from WT livers 3 and 7 dpi. Data representative of two experiments, each with 4-6 mice per genotype. ns, not significant, *p<0.05, and ****p<.0001, by Two-way ANOVA. (G) Spatial transcription of *Nos2* expression within WT granulomas at indicated timepoints.

*Nos2*^*–/–*^ mice had greater areas of the liver affected, and their granulomas were markedly abnormal (Figure 6E). Granulomas in *Nos2*^−/−^ mice did have a necrotic core surrounded by a coagulative necrotic zone. However, in contrast to WT mice, beyond the coagulative necrotic hepatocytes we observed multiple zones of Ly6G-positive necrosis with interspersed additional fragmented coagulative necrotic zones. *C. violaceum* staining appears lighter compared to WT granulomas, however, closer examination revealed that the bacteria were more dispersed throughout the granuloma and present in more discrete clusters akin to a 1 dpi lesion in WT mice (Figure 6E and 1I). This suggested that *C. violaceum* breached the granuloma and initiated new neutrophil swarms. Indeed, we observed *C. violaceum* staining throughout the zones in *Nos2*^−/−^ mice even reaching the edge of the granuloma.

The macrophage zone abnormalities in *Nos2*^−/−^ mice varied between granulomas. Some *Nos2*^*–/–*^ granulomas had single breaches in the macrophage zone (shown below in Figure 7), whereas other infected granulomas lost the macrophage zone altogether (Figure 6E). This loss of architecture could be due to the large influx of neutrophils that have overrun the granuloma response, potentially disrupting nascent macrophage zones. Neutrophil and macrophage numbers in the liver were equal by flow cytometry at 3 dpi in WT and *Nos2*^−/−^ mice, but at 7 dpi we observed significantly more neutrophils and fewer macrophages in *Nos2*^*–/–*^ mice (Figure 6F). This is consistent with histological staining for neutrophil and macrophage markers (Figure 6E).

**Figure 7.**
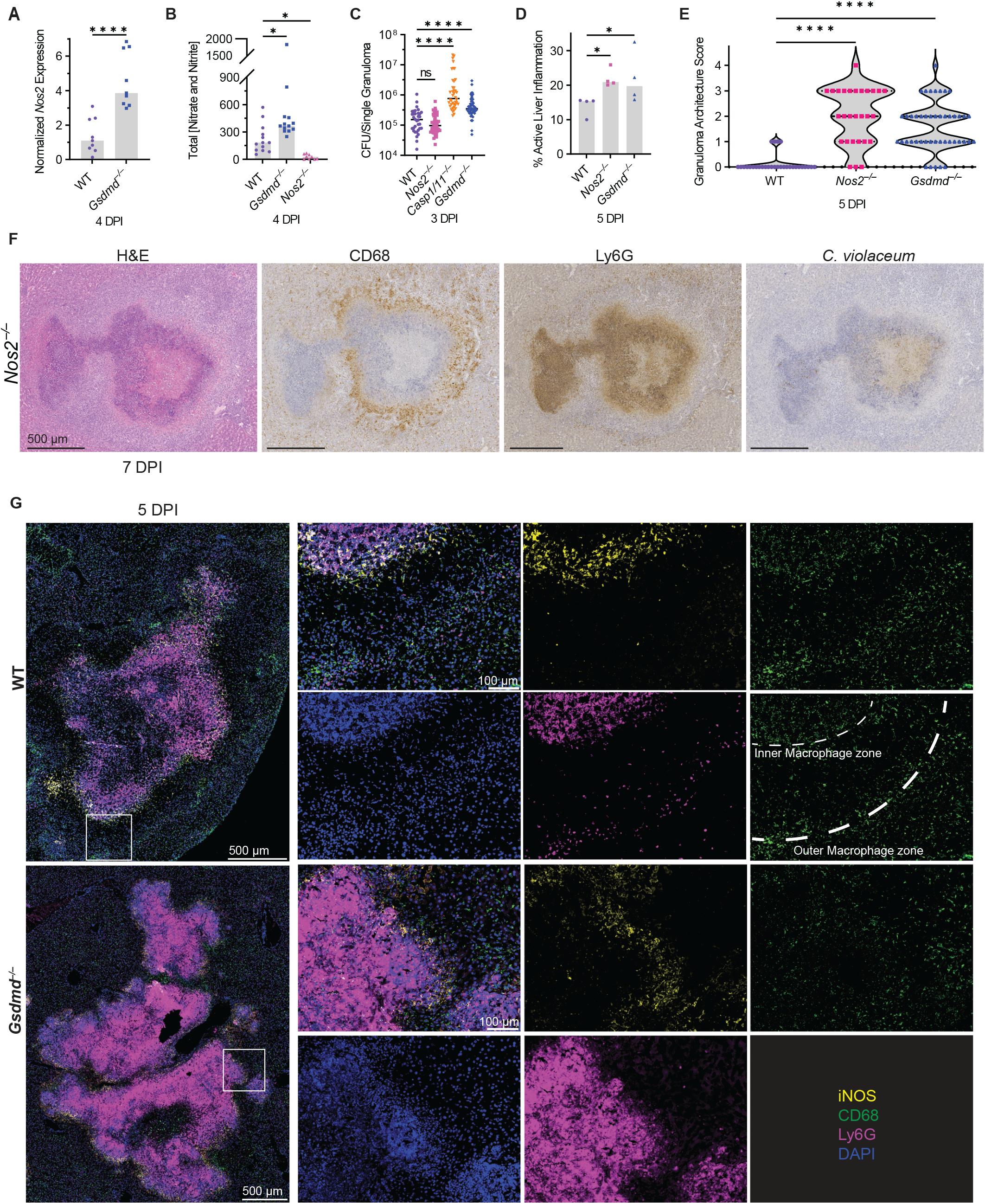
Gasdermin D and iNOS are independently required to maintain the granuloma architecture. (A-G) Mice were infected with WT *C. violaceum*. (A) qPCR of *Nos2* gene expression in WT granulomas 4 dpi. Representative of 2 experiments, each point is 5-6 pooled granulomas harvested from single mouse. ****p<.0001, by Mann-Whitney test. (B) Total nitrate and nitrite concentrations of serum from WT mice 4 dpi. Representative of 2 independent experiments performed in triplicate. *p<0.05, by Kruskal-Wallis test. (C) Bacterial burdens of single granulomas at 3 dpi. Data combined from 2 experiments, each point is a single granuloma. Solid line, data median. ****p<.0001, by One-way ANOVA test. (D) Percent active inflammation quantified per whole liver section stained with H&E. Representative of 2 experiments, each point is a single mouse. *p<0.05, by Kruskal-Wallis test. (E) Granulomas scored for failure of granuloma architecture at 5 dpi. Data combined from 2 experiments, each with 3-4 mice per genotype, each with multiple granulomas per section. ****p<.0001, by Kruskal-Wallis test. (F) Serial sections stained by H&E or indicated IHC markers of a failed granuloma 7 dpi. Representative of 2 experiments, each 3-4 mice, each with multiple granulomas per section. (G) WT and *Gsdmd*^*–/–*^ liver sections stained with indicated IF markers 5 dpi. Representative of 2 experiments; each with 4 mice, each with multiple granulomas per section.

We used spatial transcriptomics to visualize the general location of *Nos2* expression in the granuloma architecture of WT mice over time (Figure 6G). As early as 12 hpi we observed modest *Nos2* expression, which became strongest at 3 dpi and continued throughout the infection (Figure 6G). This expression pattern correlated with elevated bacterial burdens in *Nos2*^−/−^ mice but was offset by 2 days (Figure 6B), at 3 dpi macrophages are maximizing *Nos2* mRNA, but bacterial burdens in *Nos2*^−/−^ mice are not elevated until 5 dpi (Figure 6C). To reconcile this discrepancy, we stained for iNOS protein. Despite spatial transcriptomics detection of *Nos2* mRNA as early as 12 and 24 hpi, iNOS staining was not detectable at 1 dpi, and staining at 3 dpi was quite faint (Figure S5E). iNOS staining became prominent only at 5 dpi (Figure S5F). Thus, *Nos2* mRNA expression precedes detectable iNOS protein content as well as bacterial clearance. Altogether, this data demonstrates that granuloma macrophages express iNOS, that NO can kill *C. violaceum*, and in the absece of iNOS the granuloma cannot maintain its organized architecture.

### Gasdermin D and iNOS are independently required to maintain the granuloma architecture

Both *Gsdmd*^−/−^ and *Nos2*^−/−^ mice have disrupted granuloma architecture and fail to clear *C. violaceum*. We examined whether iNOS is downstream of gasdermin D in this granuloma response. This was not the case, since *Nos2* expression and NO production in *Gsdmd*^*–/–*^ mice were elevated compared to WT mice (Figure 7A and 7B). Furthermore, burdens per granuloma were elevated in *Casp1/11*^−/−^ and *Gsdmd*^−/−^ mice at 3 dpi, but not in *Nos2*^−/−^ mice (Figure 7C). Whereas *Casp1/11*^−/−^ and *Gsdmd*^−/−^ mice support splenic *C. violaceum* replication at 3 dpi, *Nos2*^−/−^ have little to no spleen burdens at this timepoint (Figure 4B, 4C, and 6B). To quantitate the defect in the overall granuloma response, we measured the area of total active inflammation in the liver section as a percentage of the total tissue area. Both *Casp1/11*^*–/–*^ and *Gsdmd*^*–/–*^ mice had increased inflammation at 3 dpi (Figure S5G and S5H), which correlates with increased bacterial burdens (Figure 4B, 4C). At this same timepoint, *Nos2*^−/−^ mice did not have increased inflammation or burdens (Figure S5G, S5H, and 6B). It was not until 5 dpi when both *Gsdmd*^*–/–*^ and *Nos2*^*–/–*^ mice had increased inflammation (Figure 7D).

Defective granulomas are heterogeneous. To quantitate the severity of failed granuloma architecture, a board-certified pathologist blindly scored each granuloma for the presence of granuloma zones, containment of Ly6G-positive cells, and the integrity of the macrophage zone (Figure 7E). A score of zero indicates fully intact granuloma architecture (as in Figure 1C). An intermediate score includes granulomas with discreet breaches in the macrophage zone. Such breaches are typically connected to a budding morphology that consists of a neutrophil swarm with *C. violaceum* staining that appears similar to a 1 dpi lesion (Figure 7F compared to Figure 1I). A score of four represents a complete loss of architecture (e.g. *Gsdmd*^−/−^ in Figure 4G and *Nos2*^−/−^ in Figure 6E). Both *Gsdmd*^−/−^ and *Nos2*^*–/–*^ mice had a markedly increased number of failed granulomas compared to WT mice (Figure 7E). These data indicate gasdermin D and iNOS are both independently required to maintenance of granuloma architecture.

We further examined *Nos2* expression in the granuloma. *Nos2* expression was associated with spatial transcriptomic clusters that contain CD45-expressing immune cells (Figure S5I and 5C); the highest level of *Nos2* expression were in clusters 6, 8, 9, and 11 (Figure S5I). These clusters with high *Nos2* expression were located at the periphery of the necrotic core and within the coagulative necrotic zone, and surprisingly less associated with the macrophage zone (Figure 5E). We had previously only noted a single macrophage zone at the periphery of the granuloma. However, instructed by these *Nos2* spatial expression patterns, we reevaluated the *C. violaceum* granuloma histology focusing on the border between the necrotic core and the coagulative necrotic hepatocytes. Indeed, in addition to the “peripheral macrophage zone” we observed a second, albeit thinner, “inner macrophage zone” between the coagulative necrotic hepatocytes and the necrotic core (Figure 7G and 2C). Therefore, in addition to validating gene candidates, spatial genomics highlighted new granuloma zones that were less apparent by H&E staining.

We next stained for iNOS protein to determine the relative expression in these two macrophage zones. Consistent with the spatial transcriptomics data, iNOS overlapped with CD68 staining in the inner macrophage zone (Figure 7G and S5F). *Gsdmd*^*–/–*^ mice have lost their granuloma architecture but retain CD68 staining, albeit discontinuously, at the periphery of the granuloma. Concomitantly, iNOS staining was maintained at the periphery of the granuloma, and was also weakly visible within hepatocytes based on nuclei staining and lack of CD68 and Ly6G staining (Figure 7G and S5F). Hepatocytes are known to produce NO via iNOS under stressful conditions^55,56^, which could occur due to the failed granuloma response *Gsdmd*^−/−^ mice. However, the NO produced by hepatocytes is not enough to rescue the failed granuloma response in these knock out mice. We again observed that *Gsdmd*^−/−^ and *Nos2*^−/−^ mice had *C. violaceum* staining throughout the entire granuloma, not limited to the core (Figure S5J).

Therefore, both gasdermin D and iNOS are required to keep *C. violaceum* contained in the granuloma core. The *C. violaceum*-induced granuloma requires at least two separate defense pathways, gasdermin D and iNOS, to maintain the integrity of the granuloma architecture, which is essential to eradicate *C. violaceum* infection.

## Discussion

*C. violaceum* first infects hepatocytes and rapidly leads to a neutrophil swarm within 1 day. Remarkably, *C. violaceum* continues to replicate through day 3 despite the copious numbers of neutrophils. This is a catastrophic failure of the innate immune response. We speculate that this failure of neutrophils to kill the bacteria is the primary problem that triggers the granuloma response. *C. violaceum* has numerous virulence traits, including T3SS effectors, most of whose functions remain unknown^2^, which could cause this neutrophil failure. In other infectious models, such as *Yersinia* species, the same neutrophil failure could arise from virulence traits that mitigate neutrophil function^57^. Indeed, pyogranulomas form in response to *Yersinia pseudotuberculosis* infection in the small intestine, which is driven by monocytes^12,58-61^. We speculate that during *C. violaceum* and *Y. pseudotuberculosis* infections it is the failure of neutrophils that triggers the granuloma response. Neutrophils have also been recently implicated in the granuloma response to *M. tuberculosis*, with recent studies showing that *M. tuberculosis* preferentially occupies and replicates in live or necrotic neutrophils^62-64^. However, this occurs once the granuloma is already formed, and other studies show neutrophils are not the first immune cell to be infected^65,66^. Thus, whether neutrophils play a key role in granuloma initiation remains unclear in the *M. tuberculosis*-induced granuloma. In contrast, by providing well-defined early events that precede granuloma formation, we propose the *C. violaceum*-induced granuloma may be revealing a fundamental reason that would explain granuloma formation.

In studies of *M. tuberculosis* infection in mice, pyroptosis-deficient mice sometimes have exacerbated infections and a failed granuloma response^27^, whereas others show no role^28,29^. This is in contrast to the clear and robust protective role of pyroptotic proteins in the *C. violaceum*-induced granuloma. Although *C. violaceum* can inhibit the apoptotic caspases^6,67^, it does not inhibit the pyroptotic caspases. We demonstrated that caspase-1/11 and gasdermin D-mediated defenses are essential to maintain granuloma architecture. Without these pyroptotic proteins, mice still express *Nos2* and generate NO, but this antibacterial mechanism fails clear the infection. During *C. violaceum* infection, WT mice have less total liver inflammation from the granuloma response compared to *Casp1/11*^−/−^ and *Gsdmd*^−/−^ mice, and these pyroptosis proteins prevent the spread of the bacteria to the edge of the granuloma. We speculate that pyroptosis prevents *C. violaceum* from establishing a replicative niche within the organized granuloma macrophage zones. Thus, we do not see a role for the pyroptotic proteins in driving a pathological exacerbation of the granuloma inflammatory response, and instead we demonstrate a clear beneficial role during the successful granuloma response.

Many other granuloma models have had contrasting conclusions as to whether caspase-1/11 and gasdermin D contributed to granuloma biology. During *Schistosomiasis*, where granuloma formation is pathological, multiple pyroptotic pathway knockout mice have smaller granulomas^68-70^, suggesting that pyroptotic proteins are detrimental. This conclusion is the opposite to our conclusion with *C. violaceum*, and may be due to the fact that schistosomes are large extracellular parasites, thus pyroptosis is not needed to prevent intracellular infection.

Similarly, in a noninfectious granuloma response, mice treated with trehalose 6.6’-dimycolate, a mycobacterial glycolipid, *Nlrp3*^*–/–*^ mice had fewer granulomas compared to WT mice^71^. Thus, in models where granulomas drive pathological tissue damage, inflammasomes can exacerbate this.

We show that iNOS is absolutely essential for the granuloma response to *C. violaceum*. WT mice use iNOS to eradicate *C. violaceum* and thus they survive; in contrast, *Nos2*^−/−^ mice succumb to the infection due to granuloma failure. *Nos2*^−/−^ mice also have increased susceptibility to *M. tuberculosis* infection^62^, however, unlike *C. violaceum* infection, iNOS fails to clear *M. tuberculosis* in mice and WT mice fail clear *M. tuberculosis* infection^72,73^. We speculate that *M. tuberculosis* has strong NO resistance mechanisms, thwarts the sterilizing effects of NO, and thereby is able to survive in granulomas long term^72^. During *M. tuberculosis* infection, NO also plays an immunomodulatory role by suppressing IL-1 production to limit pathology and prevent neutrophil recruitment^49,50^. If we are correct that *M. tuberculosis* is highly NO resistant, it may be that the primary role of NO during *M. tuberculosis* infection is to modulate the inflammatory response. We also observe increased neutrophil recruitment and a loss of granuloma architecture in *Nos2*^*–/–*^ mice, however, if *C. violaceum* is sensitive to NO killing, this cannot be separated from immunomodulation. We speculate that during *C. violaceum*-induced granuloma response the primary role of NO is to kill bacteria, and impeding neutrophil influx is a secondary function that may or may not be essential. There is another key difference between iNOS function in these two granuloma-inducing pathogens – during *M. tuberculosis* infection, *Nos2* expression is driven by the adaptive immune response^49^, whereas during *C. violaceum* infection the adaptive immune response is dispensable. Thus, the *C. violaceum*-induced granuloma demonstrates that innate immunity can use iNOS to sterilize a granuloma without adaptive T cell help.

The current understanding of granuloma biology remains perplexing, despite being studied for over a century in various granuloma models^7,14^. Granuloma biology is intriguing due to the sophisticated organization of immune cells unified into a multicellular collective to combat a pathogen that poses a dire threat to the host. Understanding how a granuloma operates requires animal model systems where the complexities of the immune response can be studied in its entirety, from the very first hours of infection to the final stages of resolution. *C. violaceum* provides such a model, where a granuloma rapidly forms and efficiently eradicates this environmental pathogen. That the granuloma successfully clears *C. violaceum* is remarkable because many pathogens survive within granulomas chronically. It may be that such host-adapted pathogens have virulence factors that thwart the efficacy of the granuloma, whereas environmental pathogens such as *C. violaceum* lack this capacity. Therefore, the *C. violaceum*-induced granuloma model could be a new way to understand the complexities of granuloma biology and clearly demonstrates that granulomas are potent immune defenses that can successfully eradicate infection.

## Acknowledgments

We thank R. Flavell and R. Vance for sharing mice directly, or through The Jackson Laboratory. This work was supported by the following NIH grants: AI133236, AI139304, AI136920 (E.A.M.) AI133236-04S1(C.K.H); NSF Graduate Research Fellowship Program DGE 2139754 (T.J.A.).. We thank the Histology Research Core Facility in the Department of Cell Biology and Physiology at the University of North Carolina, Chapel Hill NC for all paraffin embedded histological services. We thank the Molecular Genomics Core at Duke University for processing and analyzing the spatial transcriptomics (10X Genomics Visium). We thank F. Lakhani, M. Artunduaga, S. Pereles, A. Vaidyanathan, R. Eplett, M. Mann, S. Redecke, L. Scarpelli, A. Bryan, and F. Lin for mouse colony upkeep.

## Author contributions

C.K.H. performed most of the experiments with T.J.A., M.A.D., C.A.L., V.I.M., and Z.P.B. performing some experiments. E.A.M. supervised the overall project. S.A.M. supervised histology analysis. H.N.L. and B.D.M. managed the mouse colony. C.Y. and C.J.B performed the module-based analysis of spatial sequencing data; supervised by D.R.S. M.E.A performed flow cytometry and analysis. C.K.H. and E.A.M. wrote the paper.

## Competing interests

C.A.L. is employed by AbbVie. This article is composed of the authors’ work and ideas and does not reflect the ideas of AbbVie.

## METHODS

### Bacterial Strains and Culture Conditions

*Chromobacterium violaceum* ATCC 12472 was used in all experiments except mixed inocula where its nalidixic acid spontaneous resistant mutant CVN was used as WT and was mixed with CVN Δ*vioA* strain 1:1 for infection. All strains were grown on a brain heart infusion (BHI)a agar plates overnight at 37°C. Bacteria were subcultured in 2 mL BHI broth with aeration at 37°C overnight and directly diluted to 1 × 10^4^ CFU/ml PBS for infection inoculum.

### Mouse Infection and Survival

For all experiments 8- to 12-week-old mice were infected via intraperitoneal route at indicated CFUs of bacteria in PBS. Mice were monitored for survival or whole livers, spleens, and individual granulomas were harvested at indicated time points and homogenized and plated on BHI and grown as described above.

### Mouse Strains

All mouse strains were bred and housed at Duke University in a pathogen-specific free facility. For infection mice were transferred to a BSL2 infection facility within Duke University, and mice allowed to acclimate for at least two days before infection. Wild type C57BL/6 (Jackson Laboratory), *Ncf1*^*mt/mt*^ (referred to as *Ncf1*^*–/–*^; Jackson #004742), *Casp1*^*–/–*^*Casp11*^*129mt/129mt*^ (referred to as *Casp1/11*^*–/–*^)^75^, *Gsdmd*^*–/–* 76^, *Rag1*^*–/–*^(Jackson #0022216), *Prf1*^*–/–*^ (Jackson #002407), *Casp7*^*–/–*^ (Jackson #006237), *Nos2*^*–/–*^ (Jackson #002609) and *Acod1*^*–/–*^ (Jackson #029340) mice. Animal protocols were approved by the Institutional Animal Care and Use Committee (IACUC) at the University of North Carolina at Chapel Hill or by the IACUC at Duke University and met guidelines of the US National Institutes of Health for the humane care of animals.

### Histology

For paraffin embedded tissues, whole livers were harvested from euthanized mice and placed in 10% buffered formalin (VWR Cat. No. 16004-128) in 50 mL conical tubes. Tubes were inverted and swirled every other day for a minimum of 3 days to allow for full penetration of formalin into the tissue. Once fixed the liver were placed in tissue cassettes and transferred to the Histology Research Core at the University of North Carolina at Chapel Hill for embedding, cutting, slide mounting, and staining. The Histology Research Core performed all H&E and Sirius Red staining and serial sectioning of paraffin embedded tissues. For frozen fixed tissues, mice were whole body perfused through the heart with 2% paraformaldehyde (diluted down in PBS from 16%; VWR Cat. No. 15710-S), then livers were harvested and placed in 2% paraformaldehyde for 24 hours at 4°C then placed in 30% sucrose (Sigma-Aldrich Cat. No. S1888) in PBS for 48 hours at 4°C. After incubating in 30% sucrose livers were placed in OCT media (Sakura Cat. No. 4583) at room temperature for 4 hours then inserted into frozen tissue cassettes filled with OCT media and frozen at -80°C. Frozen tissues were sectioned in our lab using a cryostat (Cryostar NX70). Pathologic analysis was performed with oversight from a board-certified veterinary pathologist (S.A.M).

### Tissue Processing for Bacterial Plating

Whole livers were harvested at indicated timepoints and placed into a 7 ml homogenizer tube (Omni International Cat. No. 19-651) containing 3 mL sterile PBS and one 5 mm stainless steel bead (QIAGEN Cat. No. 69989). Spleens and single granulomas were harvested at indicated timepoints and placed into a 1 ml homogenizer tube (Fisher Brand Cat. No. 14-666-315) containing 1 mL sterile PBS and one 5 mm stainless steel bead. All were homogenized on a Fisher Brand Mead mill 24 homogenizer for livers and a Retsch MM400 homogenizer for spleens and single granulomas. After homogenization, tissue lysates were serially diluted 1:5 in sterile PBS and plated on BHI agar plates. Plates were incubated overnight at 37°C and colony forming units counted.

### Immunohistochemistry and Immunofluorescent Staining

For immunohistochemistry of paraffin embedded tissue, slides with 5 µm thick tissue sections were washed in xylene (Epredia Cat. No. 9990501) for 15 minutes, then 100% ethanol (Sigma-Aldrich Cat. No. E7023) 3 times for 3 minutes each, then 95% ethanol two times for 3 minutes each, then 80% ethanol for 3 minutes, then washed in ddiH2O for 8 minutes to rehydrate the tissues. For antigen retrieval slides were incubated in sodium citrate buffer (2.94 g Tri-Sodium Citrate, dH20 1 L, mix to dissolve, pH to 6.0 with 1N HCl, add 0.05 mL Tween 20) and pressure cooked in a pressure cooker (Instant Pot Ultra) on the high pressure setting for 12 minutes. Slides were cooled for a minimum of 20 minutes in a slide rack at room temperature and then washed in TBS-T 1x (100mL 10xTBS, 900mL ddiH20, 1mL Tween-20) for 1 minute. A pap pen was used to make a hydrophobic boarder around the tissues and then tissues were blocked with normal goat serum blocking solution, 2.5% (Vector Laboratories S-1012-50) at room temperature for 30 minutes. Tissues were washed with TBS-T 1X, then tissue was blocked with an Avidin/Biotin Blocking Kit (Vector Laboratories SP-2001) following manufactures protocol, then washed in TBS-T 1X again. Primary antibodies IHC: *C. violaceum*, 1:2000 (rabbit antisera custom generated by Cocalico Biologicals); Ly6G, 1:300 (Biolegend Cat. No. 127601); CD68, 1:200 (Abcam Cat. No. ab125212); CD3, 1:500 (Abcam Cat. No. ab5690); GFP, 1:200 (Invitrogen Cat. No. A11122); were diluted in SignalStain antibody diluent (Cell Signaling 8112L) and incubated overnight at 4°C in a humidity chamber. Then slides were washed with TBS-T 1X for 1 minute three times, then blocked for 10 minutes with 3% hydrogen peroxide (Sigma Cat. No. H1009) to block endogenous peroxidase activity. Slides were washed 3 times for 1 minute then incubated with a secondary antibody polymer (Cell Signaling-Signal Stain boost, HRP anti-rabbit, 8114; ImmPRESS® HRP Goat Anti-Rat IgG Polymer Detection Kit, Peroxidase, Cat. No. MP-7404-50). Slides were washed 3 times for 5 mintues, then stained with a DAB Substrate Kit, Peroxidase (HRP), (3,3’-diaminobenzidine; Vector Labs SK-4100) for 30-45 seconds. Then counter stained with Harris Modified Hematoxylin (Epredia Cat. No. 72704) or Meyers Modified Hematoxylin (Abcam Cat. No. ab220365) for 10-30 seconds. Hematoxylin was blued using running tap water for 1 minute. Slides were dehydrated in 50% ethanol for 2 minutes, then 70% ethanol for 2 minutes, then two times in 100% ethanol for 2 minutes, then two times xylene for 5 minutes. Slides were then covered with Permount mounting medium (Fisher Chemical SP15-100) and covered with a coverslip.

For Immunofluorescence on frozen tissue (5 µm thick tissue sections), slides were brought to room temperature for 15-20 minutes and a pap pen was used to draw a hydrophobic barrier around the tissue. Tissues were blocked with 2% Fc block (BD Pharmingen Cat. No. 553142) with 5% normal goat serum in 1X PBS at room temperature for one hour. Slides were washed for 1 minute in 1X PBS and then incubated with primary antibody overnight at 4°C in a humidity chamber. Primary antibodies IF: Ly6G Alexa Fluor 647, 1:100 (Biolegend Cat. No. 127610), CD68 Alexa Fluor 488, 1:100 (Abcam Cat. No. ab201844), iNOS Alexa Fluor 568, 1:100 (Abcam Cat. No. ab209595), e-cadherin, 1:200 (Abcam Cat. No. ab15148). Slides were then washed in 1X PBS three times for 5 minutes and incubated with a secondary antibody (Invitrogen Goat-anti-rabbit 594, 1:1000, Cat. No. A32740) at room temperature for an hour. Slides were washed in 1X PBS three times for 5 minutes and mounted using Fluoroshield™ with DAPI (Sigma-Aldrich F6057) and covered with a coverslip.

### Histology Image Capture

All histology images were captured using a Keyence all-in-one Fluorescent Microscope BZ-X800/BZX810. For histology image stitching and IF analysis the BZ-X800 analyzer was used.

### 10x Genomics Visium

Infected mouse liver tissue was harvested at various time points and embedded in OCT (Sakura Cat. No. 4583) and frozen. Frozen livers were cut to optimal section thickness and placed on a Tissue Optimization Slide to determine permeabilization conditions. After, tissues were then placed within a 6.5mm^2^ field on an expression slide that contains 5,000 barcoded probes. The tissues were then fixed and stained with Hoechst and Eosin then permeabilized to release mRNA which binds to spatially barcoded capture probes, allowing for the capture of gene expression information. Captured mRNA from the slide surface was denatured, cleaved, and transferred into a PCR tube. From there, the cDNA was amplified and standard NGS libraries were prepared. Adapters were ligated to each fragment followed by a sample index PCR. The libraries were sequenced to an average of 50,000 reads/probe on a paired end, dual indexed flowcell in the format of 28×10×10×90. Data was then uploaded to analysis packages for visualization.

Visium spatial data was analyzed using 10xGenomincs Space Ranger software and visualization through Loupe Browser^77,78^. Secondary statistical analysis will be performed using an R package in Seurat^79^. Linear dimensional reduction was preformed to calculate principal components using the most variably expressed genes in the spatial data. Spots were grouped into an optimal number of clusters for de novo cell type discovery using Seurat’s FindNeighbors() and FindClusters() functions, graph-based clustering approaches with visualization of spots is achieved through the use of manifold learning technique UMAP (Uniform Manifold Approximation and Projection)^80^. SpatialDimPlot() function in Seurat overlays the clustering results on the image to give combined plot of the expression data and the histology image. Additional downstream analyses include examining the distribution of a priori genes of interest, closer examination of genes associated with spot clusters, and the refined clustering of spots in order to identify further resolution of cell types, in addition to comparing differences between experiments of different states.

### Module-Based Analysis of Spatial Sequencing Data

To understand the cellular composition of each cluster, we leveraged two published single cell RNA-seq datasets^46,47^ from mouse liver and utilized the markers for immune cells and non-hematopoietic cells to calculate the module scores of different major cell types (Table S4E). The module score was generated as previously described^81^ using the AddModuleScore function from the Seurat R package version 4^82^. Results were visualized using the ggplot2 R package and used to track the approximate relative abundance of unique cell types or specific genes at each infection time point. Abbrev. Key for our reference only: NC = Necrotic Core, CN = Coagulative Necrosis, M = Macrophage, EC = Endothelial Cell, OG = Outside Granuloma, Hep = Hepatocyte

### In vitro killing of *C. violaceum* with Itaconate and Nitric Oxide

One WT *C. violaceum* colony was grown overnight in BHI broth then diluted to 1×10^6^ CFU/mL of PBS and mixed with various concentrations of DEA NONOate (Cayman Chemical Cat. No. 82100) with or without itaconic acid (Sigma-Aldrich Cat. No. I29204). Cultures were incubated at room temperature for 3 hours then diluted at 1:5 in sterile PBS and plated on BHI agar plates to assess bacterial viability.

### Flow Cytometry

Whole livers were harvested at the indicated time points post infection. Briefly, mice were euthanized according to IACUC guidelines, followed by whole body perfusion with PBS (Gibco, Cat. No. 14190-144). Whole livers were harvested and minced on ice using scissors, followed by incubation in digestion buffer (100U/mL Collagenase Type IV (Gibco, Cat. No. 17104019) prepared in DMEM (Gibco, Cat. No. 11885-084), supplemented with 1 mM CaCl_2_ and 1 mM MgCl_2_,) at 37°C for 40 minutes with intermittent vortexing. Digested tissues were mechanically homogenized through a Falcon® 40 µm cell strainer (Corning, Cat. No. 352340) to remove the majority of hepatocytes, and washed twice with RPMI (Gibco™, Cat. No. 11875-093) supplemented with 1% FBS (CPS Serum, Cat. No. FBS-500-HI) and 1X penicillin/streptomycin (Gibco, Cat. No. 15140-122), followed by centrifugation in an Eppendorf® centrifuge (model 5810 R) at 1200 rpm (290g) for 8 minutes at room temperature. Leukocytes were further isolated using a Percoll® gradient: samples were resuspended in 45% Percoll® (GE Healthcare, Cat. No. 17-0891-01), prepared in DMEM + 1.5M NaCl), with an 80% Percoll® (prepared in PBS + 1.5 M NaCl) underlay, and spun for 20 minutes at 2000 rpm (805g), room temperature, with no brake. Following collection of the leukocyte layer at the gradient interface, samples were washed twice, as before, and red blood cells were lysed with 1X RBC Lysis Buffer (eBioscience, Cat. No. 00-4333-57, according to product manual). Cells were washed and counted using trypan blue. 1×10^6^ cells from each liver were stained for various cell markers: Live-or-Dye™ fixable viability dye in APC-Cy™7 (Biotium, Cat. No. 32008, according to product manual), Mouse BD Fc Block™ (BD Biosciences, Cat. No. 553142, according to product manual) rat anti-mouse Ly-6G in BV421™ at 1:300 for 30 minutes (BD Horizon™, Cat. No. 562737), and finally, rat anti-mouse CD68 in FITC at 1:300 for 30 minutes (BioLegend®, Cat. No. 137005) using Intracellular Fixation & Permeabilization Buffer (eBioscience, Cat. No. 88-8824-00, according to product manual). Cells were acquired on a BD LSRFortessa X-20 Cell Analyzer (Duke Flow Cytometry Core Facility), and analyzed using FlowJo (for Windows, version 10.7.1).

### Quantification of relative *Nos2* expression

Granulomas were excised from infected livers and placed into 1 ml homogenizer tube (Fisher Brand Cat. No. 14-666-315) containing 1 mL sterile PBS and one 5 mm stainless steel bead (QIAGEN Cat. No. 69989). Nucleic acids were isolated from excised granulomas by homogenizing in TRIzol (Invitrogen, Cat. No. 15596026) for 10 minutes. After homogenization, RNA was isolated by adding chloroform to a final ratio of 1:5 chloroform:TRIzol, centrifuging, then mixing the aqueous layer with 70% EtOH, then applying to Qiagen RNeasy columns to isolate RNAs longer than 200 bp. RNA purification by RNeasy column was performed according to the product manual. RNA quality was evaluated by 1% bleach gel^83^. cDNA was generated from 1 ug of RNA using Oligo dT(12-18) primers (Invitrogen, 18418012) and SuperScript II reverse transcriptase (Invitrogen, 18064022). Resultant cDNA at 1:100 dilution was subject to qPCR performed using iQ SYBR Green Supermix (BioRad, Cat. No. 1708880) on a QuantStudio 3 Real-Time PCR System (Applied Biosystems, A28567). Primers for *Nos2* and *B2m* were designed using NCBI PrimerBlast^84^. *Nos2* primers were designed to span exons 12 and 13 of *Nos2*, where the calmodulin binding domain was replaced with a neomycin cassette to generate the *Nos2* knockout mice^85^. *Nos2* was amplified using primers NOS2_F and NOS2_R, and *B2m* was amplified using primers B2M_F and B2M_R. *Nos2*^*–/–*^ mice did not amplify any product with primers NOS2_F and NOS2_R by agarose gel and melt curve analysis. Quantification of relative expression of NOS2 to B2M was determined using the ΔΔCt method^86^.999 At least 3 biological replicates were sampled for each treatment, and 3 technical replicates were run for each condition.

### Primers

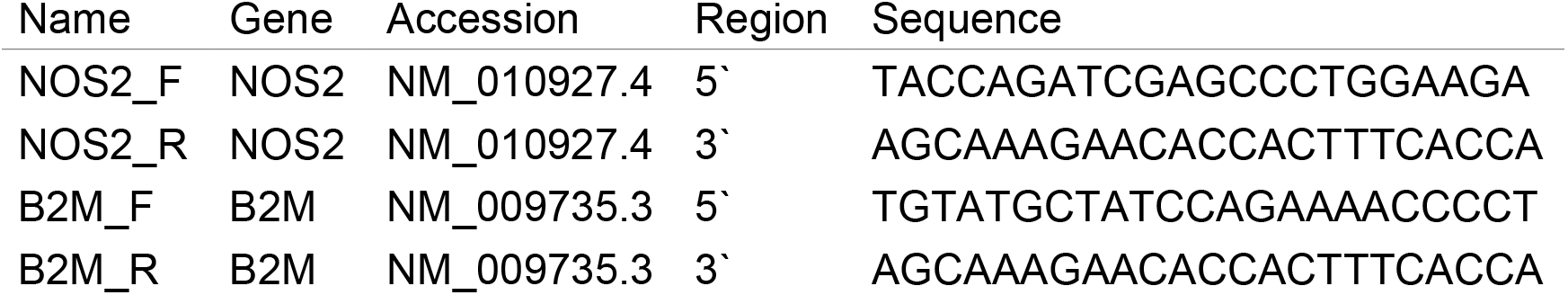

### Total Nitrate and Nitrite Assay of *C. violaceum* granulomas

Blood from indicated mouse strains was harvested via heart puncture and centrifuged at 10,000 g for 6 minutes in serum separation tubes (BD Microtainer Blood Collection Tubes Cat. No. 365967). Serum was filtered through a Microcon-10kDa Centrifugal Filter Unit with Ultracel-10 membrane (Millipore Cat. No. MRCPRT010) per company protocol. Filtered serum was diluted 1:8 in assay buffer and measured for total nitrate and nitrite using Nitrate/Nitrite Colorimetric Assay Kit (Cayman Chemical Cat. No. 780001). Absorbance was measured at 540 nm using a BioTek Synergy H1 Microplate Reader, and total nitrate and nitrite calculated from absorbance using kit protocol.

### Total Percent Inflammation Measurements

2X stitched images of whole liver were taken using a Keyence Microscope (BZ-X800). A control measurement was taken at this time using the internal Keyence measurement analysis tool to determine um^2 of control square. Images were then loaded into Microsoft Windows 10 Paint (version21H2) to trace both the whole liver and regions of inflammation. Traced images were loaded into Image J (win-64 version 1.53) and area measured using the tracing tool. Image J areas were converted into true um^2 area using the control measurement values. Percent inflammation determined by dividing the total area of inflammation measured by the total liver area measured.

### Statistical Analysis

All statistical analysis was performed with GraphPad Prism 9. Discrete data was first assessed for normal distribution using a Shapiro-Wilk normality test. Data with normal distribution was analyzed with either unpaired two-tail t test (2 groups) or a one-way ANOVA (3 or more groups). Discrete data that did not have a normal distribution, or ordinal data for histological scoring, was analyzed with a Mann-Whitney (2 groups) or Kruskal-Wallis (3 or more groups). Experiments with two factors were analyzed with a two-way ANOVA. Survival analyses were analyzed using a Kaplan-Meier survival curve with Bonferroni correction for multiple comparisons where appropriate.

## SUPPLEMENT FIGURES

**Supplement Figure 1. Characterization of the *C. violaceum*-induced granuloma**.

(A-F) Mice were infected with WT *C. violaceum* or indicated strains.

(A) Inoculum was a 1:1 mixture of WT and *∆vioA C. violaceum*. Schematic for harvesting single granulomas using circular tipped tweezers. BHI agar plate representative of 5 dpi single granulomas’ plated CFUs.

(A) Visualization of *C. violaceum* within liver Kupffer cells and infiltrating immune cells 12 hpi. Representative of 2 experiments, each with 3-4 mice.

(B) Zoomed in CD68 staining of 1 dpi lesion from Figure 1I.

(C) Zoomed in IF staining of 5 dpi granuloma of indicated markers. Representative of 2 experiments, each with 3-4 mice, each with multiple granulomas per section.

(D) H&E staining of resolving granulomas with no necrotic core 14 dpi from Figure 3A.

(E) WT liver sections stained with indicated IF markers 14 dpi. Representative of 2 experiments; each with 4 mice, each with multiple granulomas per section.

**Supplement Figure 2. Kinetics of granuloma resolution**.

(A-G) Mice were infected with WT *C. violaceum*.

(A) Bacterial burdens in the liver at indicated dpi. Data combined from 2 experiments, each point is a single mouse.

(B) Bacterial burdens from (A) displayed as % bacterial clearance.

(C) Serial sections stained by H&E or indicated IHC markers of large granuloma from Figure 3A.

(D) Representative whole liver sections stained with H&E used for percent active inflammation quantification from Figure 3G. Whole livers for 5, 7, 14, 21 dpi are same as whole livers in Figures 2A, 3A, and 3C.

(E) Bacterial burdens from single granulomas 7 and 10 dpi. Representative of 2 experiments, each point is a single granuloma, each letter is a mouse.

(F) WT granuloma stained for CD3 at 7dpi.

(G) H&E staining of *Rag1*^*–/–*^ livers at 26 dpi. Data represents one experiment with 3 mice. Dashed line, limit of detection; solid line, median.

**Supplement Figure 3. Thrombosis, Ischemia, and other aspects of the granuloma**.

(A-G) Mice were infected with WT *C. violaceum*.

(A) Isolate ischemia stained with H&E 3 dpi. Representative of 10 experiments, each with 3-4 mice, multiple areas in section.

(B) Adjacent ischemia stained with H&E 3 dpi. Representative of 10 experiments, each with 3-4 mice, each with multiple areas in section.

(C and D) Adjacent ischemia stained with H&E and IHC for *C. violaceum* 3 dpi. Representative of 10 experiments, each with 3-4 mice, each with multiple areas in section.

(E) Clot stained with H&E 7 dpi. Arrows, fibrin strands. Representative of 5 experiments, each with 3-4 mice, each with multiple clots per section.

(F) Table of total WT survival numbers from all survivals in paper.

(G) *NCR1*^*gfp*^ mice stained for indicated IHC marker 3 dpi. Arrows, NK cells. Representative of one experiment with 3 mice.

(H) Survival analysis of *Perf*^*–/–*^ mice. Data represents one experiment with 4 mice.

(I) Bacterial burdens from liver and spleen 19 dpi from (H), each point is a single mouse. Dashed line, limit of detection; solid line data median.

(J) Representative granuloma histology of WT, *Perf*^*–/–*^, and *Casp7*^*–/–*^ mice at 3 dpi stained for H&E. Histopathology analyses were performed on livers from 4 mice per time point from one experiment.

(K) IHC staining of *C. violaceum* dissemination in liver of *Casp1*^*–/–*^*Casp11*^*–/–*^ (*Casp1/11*^*–/–*^) mice 3 dpi. Arrows, disseminate bacteria.

(L) IHC staining of *C. violaceum* dissemination in liver of *Gsdmd*^*–/–*^ mice 3 dpi. Arrows, disseminate bacteria.

**Supplement Figure 4. Spatial transcriptomics of the granuloma**.

(A-F) Mice were infected i.p. with 1 × 10^4^ CFU WT *C. violaceum* and one liver was harvested per indicated timepoint for spatial transcriptomics analysis.

(A) H&E and spatial transcriptomic orientation of indicated clusters from all time points; same granuloma at 5 dpi from Figure 5A.

(B) UMAP plot for cluster expression orientation for clusters from all timepoints of infection expressing *Ptprc*.

(C) UMAP plot for cluster expression orientation for clusters from all timepoints of infection expressing *Alb*.

(D) UMAP plot for cluster expression orientation for clusters’ location from all timepoints of infection within the granuloma architecture.

(E) Table of genes used for calculating module scores in Figure 5D.

**Supplement Figure 5. Analysis of roles for gasdermin D, iNOS, and IRG1 in the granuloma**.

(A-G) Mice were infected with WT *C. violaceum*.

(A) Bacterial burdens from *C. violaceum* incubated at indicated concentrations of DEA NONOate and Itaconic acid after 3 hours Data representative of 2 experiments, each point is a single culture. ****p<.0001, by Two-way ANOVA.

(B) Survival analysis of WT and *Acod1*^*–/–*^ mice. Data representative of 2 experiments, each with 4-5 mice per genotype. ns, not significant, by Kaplan-Meier survival analysis.

(C) Bacterial burdens from liver 7 and 10 dpi. Data combined from 2 experiments, each point is a single mouse. ns, not significant, by Mann-Whitney (7 dpi) and Unpaired two-tailed t test (10 dpi).

(D) Survival analysis of *Nos2*^*–/–*^ and littermate *Nos2*^*+/–*^ mice. Data representative of one experiment, 4 *Nos2*^*–/–*^ and 11 *Nos2*^*+/–*^ mice. ****p<.0001, by Kaplan-Meier survival analysis.

(E) WT liver sections stained with indicated IF markers 1 and 3 dpi. Representative of 2 experiments; each with 4 mice, each with multiple granulomas per section.

(F) WT, *Gsdmd*^*–/–*^, and *Nos2*^*–/–*^ mice stained with indicated IF markers 5 dpi. Same WT and *Gsdmd*^*–/–*^ representative granulomas as Figure 7G. Representative of 2 experiments, each with 3-4 mice per genotype, each with multiple granulomas per section.

(G) Percent active inflammation quantified per whole liver section stained with H&E 3 dpi. Data representative of 2 experiments, each with 3-4 mice per genotype, each point is a single mouse.

*p<0.05, by One-way ANOVA.

(H) Percent active inflammation quantified per whole liver section stained with H&E 3 dpi. Data representative of 2 experiments, each with 3-4 mice per genotype, each point is a single mouse.

*p<0.05, by One-way ANOVA.

(I) UMAP plot for cluster expression orientation for clusters from all timepoints of infection expressing *Nos2*.

(J) WT, *Gsdmd*^*–/–*^, and *Nos2*^*–/–*^ mice stained with indicated IF markers 5 dpi. Same WT and *Gsdmd*^*–/–*^ representative granulomas as Figure 7G. Representative of 2 experiments, each with 3-4 mice per genotype, each with multiple granulomas per section. Dashed line, limit of detection; solid line data median.

## Notes

https://data.mendeley.com/datasets/kmbskdyshj/draft?a=f46e6899-0c61-48b6-b6d3-ae550e0fa4c5

### Summary of Updates

We updated the figure legend for Figure 3.

## REFERENCES

1. Batista, J.H., and da Silva Neto, J.F. (2017). Chromobacterium violaceum Pathogenicity: Updates and Insights from Genome Sequencing of Novel Chromobacterium Species. Front Microbiol 8, 2213 10.3389/fmicb.2017.02213.

2. Miki, T., Akiba, K., Iguchi, M., Danbara, H., and Okada, N. (2011). The Chromobacterium violaceum type III effector CopE, a guanine nucleotide exchange factor for Rac1 and Cdc42, is involved in bacterial invasion of epithelial cells and pathogenesis. Mol Microbiol 80, 1186–1203 10.1111/j.1365-2958.2011.07637.x.

3. Maltez, V.I., Tubbs, A.L., Cook, K.D., Aachoui, Y., Falcone, E.L., Holland, S.M., Whitmire, J.K., and Miao, E.A. (2015). Inflammasomes Coordinate Pyroptosis and Natural Killer Cell Cytotoxicity to Clear Infection by a Ubiquitous Environmental Bacterium. Immunity 43, 987–997 10.1016/j.immuni.2015.10.010.

4. A M Macher, T.B.C., A S Fauci (1982). Chronic granulomatous disease of childhood and Chromobacterium violaceum infections in the southeastern United States. 97, 51–55 10.7326/0003-4819-97-1-51.

5. Nozaki, K., Maltez, V.I., Rayamajhi, M., Tubbs, A.L., Mitchell, J.E., Lacey, C.A., Harvest, C.K., Li, L., Nash, W.T., Larson, H.N., et al. (2022). Caspase-7 activates ASM to repair gasdermin and perforin pores. Nature 606, 960–967 10.1038/s41586-022-04825-8.

6. Peng, T., Tao, X., Xia, Z., Hu, S., Xue, J., Zhu, Q., Pan, X., Zhang, Q., and Li, S. (2022). Pathogen hijacks programmed cell death signaling by arginine ADPR-deacylization of caspases. Mol Cell 82, 1806–1820 e1808 10.1016/j.molcel.2022.03.010.

7. Pagan, A.J., and Ramakrishnan, L. (2018). The Formation and Function of Granulomas. Annu Rev Immunol 36, 639–665 10.1146/annurev-immunol-032712-100022.

8. Shah, K.K., Pritt, B.S., and Alexander, M.P. (2017). Histopathologic review of granulomatous inflammation. J Clin Tuberc Other Mycobact Dis 7, 1–12 10.1016/j.jctube.2017.02.001.

9. Erika Heninger, L.H.H., Jozsef Karman, Sinarack Macvilay, Bjork Hill, Jon P. Woods, and Matyas Sandor (2006). Characterization of the Histoplasma capsulatum-Induced Granuloma. J Immunol 177, 3303–3313 10.4049/jimmunol.177.5.3303.

10. Orme, I.M., and Basaraba, R.J. (2014). The formation of the granuloma in tuberculosis infection. Semin Immunol 26, 601–609 10.1016/j.smim.2014.09.009.

11. Timmermans, W.M., van Laar, J.A., van Hagen, P.M., and van Zelm, M.C. (2016). Immunopathogenesis of granulomas in chronic autoinflammatory diseases. Clin Transl Immunology 5, e118 10.1038/cti.2016.75.

12. Sorobetea, D., Matsuda, R., Peterson, S.T., Grayczyk, J.P., Rao, I., Krespan, E., Lanza, M., Assenmacher, C.-A., Beiting, D., Radaelli, E., et al. (2022). Inflammatory monocytes promote pyogranuloma formation to counteract Yersinia blockade of host defense. 10.1101/2022.02.06.479204.

13. Kaye, P.M., and Beattie, L. (2016). Lessons from other diseases: granulomatous inflammation in leishmaniasis. Semin Immunopathol 38, 249–260 10.1007/s00281-015-0548-7.

14. Ramakrishnan, L. (2012). Revisiting the role of the granuloma in tuberculosis. Nat Rev Immunol 12, 352–366 10.1038/nri3211.

15. Schwartz, C., and Fallon, P.G. (2018). Schistosoma “Eggs-Iting” the Host: Granuloma Formation and Egg Excretion. Front Immunol 9, 2492 10.3389/fimmu.2018.02492.

16. Madigan, C.A., Cameron, J., and Ramakrishnan, L. (2017). A Zebrafish Model of Mycobacterium leprae Granulomatous Infection. J Infect Dis 216, 776–779 10.1093/infdis/jix329.

17. Matyas Sandor, J.V.W., and Thomas A. Wynn (2003). Granulomas in schistosome and mycobacterial infections: a model of local immune responses. TRENDS in Immunology 24, 44–52

18. Lamps, L.W. (2015). Hepatic Granulomas: A Review With Emphasis on Infectious Causes. Arch Pathol Lab Med 139, 867–875 10.5858/arpa.2014-0123-RA.

19. Cadena, A.M., Fortune, S.M., and Flynn, J.L. (2017). Heterogeneity in tuberculosis. Nat Rev Immunol 17, 691–702 10.1038/nri.2017.69.

20. Marakalala, M.J., Raju, R.M., Sharma, K., Zhang, Y.J., Eugenin, E.A., Prideaux, B., Daudelin, I.B., Chen, P.Y., Booty, M.G., Kim, J.H., et al. (2016). Inflammatory signaling in human tuberculosis granulomas is spatially organized. Nat Med 22, 531–538 10.1038/nm.4073.

21. Egen, J.G., Rothfuchs, A.G., Feng, C.G., Winter, N., Sher, A., and Germain, R.N. (2008). Macrophage and T cell dynamics during the development and disintegration of mycobacterial granulomas. Immunity 28, 271–284 10.1016/j.immuni.2007.12.010.

22. Guirado, E., and Schlesinger, L.S. (2013). Modeling the Mycobacterium tuberculosis Granuloma - the Critical Battlefield in Host Immunity and Disease. Front Immunol 4, 98 10.3389/fimmu.2013.00098.

23. Russell, D.G. (2007). Who puts the tubercle in tuberculosis? Nat Rev Microbiol 5, 39–47 10.1038/nrmicro1538.

24. Lin, P.L., Ford, C.B., Coleman, M.T., Myers, A.J., Gawande, R., Ioerger, T., Sacchettini, J., Fortune, S.M., and Flynn, J.L. (2014). Sterilization of granulomas is common in active and latent tuberculosis despite within-host variability in bacterial killing. Nat Med 20, 75–79 10.1038/nm.3412.

25. Zhan, L., Tang, J., Sun, M., and Qin, C. (2017). Animal Models for Tuberculosis in Translational and Precision Medicine. Front Microbiol 8, 717 10.3389/fmicb.2017.00717.

26. Bucsan, A.N., Mehra, S., Khader, S.A., and Kaushal, D. (2019). The current state of animal models and genomic approaches towards identifying and validating molecular determinants of Mycobacterium tuberculosis infection and tuberculosis disease. Pathog Dis 77 10.1093/femspd/ftz037.

27. Mayer-Barber, K.D., Barber, D.L., Shenderov, K., White, S.D., Wilson, M.S., Cheever, A., Kugler, D., Hieny, S., Caspar, P., Nunez, G., et al. (2010). Caspase-1 independent IL-1beta production is critical for host resistance to mycobacterium tuberculosis and does not require TLR signaling in vivo. J Immunol 184, 3326–3330 10.4049/jimmunol.0904189.

28. McElvania Tekippe, E., Allen, I.C., Hulseberg, P.D., Sullivan, J.T., McCann, J.R., Sandor, M., Braunstein, M., and Ting, J.P. (2010). Granuloma formation and host defense in chronic Mycobacterium tuberculosis infection requires PYCARD/ASC but not NLRP3 or caspase-1. PLoS One 5, e12320 10.1371/journal.pone.0012320.

29. Dorhoi, A., Nouailles, G., Jorg, S., Hagens, K., Heinemann, E., Pradl, L., Oberbeck-Muller, D., Duque-Correa, M.A., Reece, S.T., Ruland, J., et al. (2012). Activation of the NLRP3 inflammasome by Mycobacterium tuberculosis is uncoupled from susceptibility to active tuberculosis. Eur J Immunol 42, 374–384 10.1002/eji.201141548.

30. Kauffman, K.D., Sakai, S., Lora, N.E., Namasivayam, S., Baker, P.J., Kamenyeva, O., Foreman, T.W., Nelson, C.E., Oliveira-de-Souza, D., Vinhaes, C.L., et al. (2021). PD-1 blockade exacerbates Mycobacterium tuberculosis infection in rhesus macaques. Sci Immunol 6 10.1126/sciimmunol.abf3861.

31. Chai, Q., Yu, S., Zhong, Y., Lu, Z., Qiu, C., Yu, Y., Zhang, X., Zhang, Y., Lei, Z., Qiang, L., et al. (2022). A bacterial phospholipid phosphatase inhibits host pyroptosis by hijacking ubiquitin. Science 378, eabq0132 10.1126/science.abq0132.

32. Yang, Y., Xu, P., He, P., Shi, F., Tang, Y., Guan, C., Zeng, H., Zhou, Y., Song, Q., Zhou, B., et al. (2020). Mycobacterial PPE13 activates inflammasome by interacting with the NATCH and LRR domains of NLRP3. FASEB J 34, 12820–12833 10.1096/fj.202000200RR.

33. Beckwith, K.S., Beckwith, M.S., Ullmann, S., Saetra, R.S., Kim, H., Marstad, A., Asberg, S.E., Strand, T.A., Haug, M., Niederweis, M., et al. (2020). Plasma membrane damage causes NLRP3 activation and pyroptosis during Mycobacterium tuberculosis infection. Nat Commun 11, 2270 10.1038/s41467-020-16143-6.

34. Zilu Qul, J.Z., Yidan Zhou, Yan Xie, Yanjing Jiang, Jian Wu, Zuoqin Luo, Guanghui Liu, Lei Yin, Xiao-Lian Zhang (2020). Mycobacterial EST12 activates a RACK1–NLRP3–gasdermin D pyroptosis–IL-1℘ immune pathway. Science advances 6 10.1126/sciadv.aba4733.

35. Liu, M., Zhang, S.S., Liu, D.N., Yang, Y.L., Wang, Y.H., and Du, G.H. (2021). Chrysomycin A Attenuates Neuroinflammation by Down-Regulating NLRP3/Cleaved Caspase-1 Signaling Pathway in LPS-Stimulated Mice and BV2 Cells. Int J Mol Sci 22 10.3390/ijms22136799.

36. Mishra, B.B., Moura-Alves, P., Sonawane, A., Hacohen, N., Griffiths, G., Moita, L.F., and Anes, E. (2010). Mycobacterium tuberculosis protein ESAT-6 is a potent activator of the NLRP3/ASC inflammasome. Cell Microbiol 12, 1046–1063 10.1111/j.1462-5822.2010.01450.x.

37. Saiga, H., Kitada, S., Shimada, Y., Kamiyama, N., Okuyama, M., Makino, M., Yamamoto, M., and Takeda, K. (2012). Critical role of AIM2 in Mycobacterium tuberculosis infection. Int Immunol 24, 637–644 10.1093/intimm/dxs062.

38. Rastogi, S., Ellinwood, S., Augenstreich, J., Mayer-Barber, K.D., and Briken, V. (2021). Mycobacterium tuberculosis inhibits the NLRP3 inflammasome activation via its phosphokinase PknF. PLoS Pathog 17, e1009712 10.1371/journal.ppat.1009712.

39. Kurane, T., Matsunaga, T., Ida, T., Sawada, K., Nishimura, A., Fukui, M., Umemura, M., Nakayama, M., Ohara, N., Matsumoto, S., et al. (2022). GRIM-19 is a target of mycobacterial Zn(2+) metalloprotease 1 and indispensable for NLRP3 inflammasome activation. FASEB J 36, e22096 10.1096/fj.202101074RR.

40. Master, S.S., Rampini, S.K., Davis, A.S., Keller, C., Ehlers, S., Springer, B., Timmins, G.S., Sander, P., and Deretic, V. (2008). Mycobacterium tuberculosis prevents inflammasome activation. Cell Host Microbe 3, 224–232 10.1016/j.chom.2008.03.003.

41. Subbarao, S., Sanchez-Garrido, J., Krishnan, N., Shenoy, A.R., and Robertson, B.D. (2020). Genetic and pharmacological inhibition of inflammasomes reduces the survival of Mycobacterium tuberculosis strains in macrophages. Sci Rep 10, 3709 10.1038/s41598-020-60560-y.

42. Adigun, R., Basit, H., and Murray, J. (2022). Cell Liquefactive Necrosis. In StatPearls (Treasure Island (FL))https://www.ncbi.nlm.nih.gov/pubmed/28613685

43. Miki, T., Iguchi, M., Akiba, K., Hosono, M., Sobue, T., Danbara, H., and Okada, N. (2010). Chromobacterium pathogenicity island 1 type III secretion system is a major virulence determinant for Chromobacterium violaceum-induced cell death in hepatocytes. Mol Microbiol 77, 855–872 10.1111/j.1365-2958.2010.07248.x.

44. Giacomini, E., Iona, E., Ferroni, L., Miettinen, M., Fattorini, L., Orefici, G., Julkunen, I., and Coccia, E.M. (2001). Infection of human macrophages and dendritic cells with Mycobacterium tuberculosis induces a differential cytokine gene expression that modulates T cell response. J Immunol 166, 7033–7041 10.4049/jimmunol.166.12.7033.

45. Nozaki, K., Li, L., and Miao, E.A. (2022). Innate Sensors Trigger Regulated Cell Death to Combat Intracellular Infection. Annu Rev Immunol 40, 469–498 10.1146/annurev-immunol-101320-011235.

46. Su, Q., Kim, S.Y., Adewale, F., Zhou, Y., Aldler, C., Ni, M., Wei, Y., Burczynski, M.E., Atwal, G.S., Sleeman, M.W., et al. (2021). Single-cell RNA transcriptome landscape of hepatocytes and non-parenchymal cells in healthy and NAFLD mouse liver. iScience 24, 103233 10.1016/j.isci.2021.103233.

47. Remmerie, A., Martens, L., Thone, T., Castoldi, A., Seurinck, R., Pavie, B., Roels, J., Vanneste, B., De Prijck, S., Vanhockerhout, M., et al. (2020). Osteopontin Expression Identifies a Subset of Recruited Macrophages Distinct from Kupffer Cells in the Fatty Liver. Immunity 53, 641–657 e614 10.1016/j.immuni.2020.08.004.

48. Mattila, J.T., Ojo, O.O., Kepka-Lenhart, D., Marino, S., Kim, J.H., Eum, S.Y., Via, L.E., Barry, C.E., 3rd, Klein, E., Kirschner, D.E., et al. (2013). Microenvironments in tuberculous granulomas are delineated by distinct populations of macrophage subsets and expression of nitric oxide synthase and arginase isoforms. J Immunol 191, 773–78410.4049/jimmunol.1300113.

49. Mishra, B.B., Rathinam, V.A., Martens, G.W., Martinot, A.J., Kornfeld, H., Fitzgerald, K.A., and Sassetti, C.M. (2013). Nitric oxide controls the immunopathology of tuberculosis by inhibiting NLRP3 inflammasome-dependent processing of IL-1beta. Nat Immunol 14, 52–60 10.1038/ni.2474.

50. Mishra, B.B., Lovewell, R.R., Olive, A.J., Zhang, G., Wang, W., Eugenin, E., Smith, C.M., Phuah, J.Y., Long, J.E., Dubuke, M.L., et al. (2017). Nitric oxide prevents a pathogen-permissive granulocytic inflammation during tuberculosis. Nat Microbiol 2, 17072 10.1038/nmicrobiol.2017.72.

51. Nair, S., Huynh, J.P., Lampropoulou, V., Loginicheva, E., Esaulova, E., Gounder, A.P., Boon, A.C.M., Schwarzkopf, E.A., Bradstreet, T.R., Edelson, B.T., et al. (2018). Irg1 expression in myeloid cells prevents immunopathology during M. tuberculosis infection. J Exp Med 215, 1035–1045 10.1084/jem.20180118.

52. Bomfim, C.C.B., Fisher, L., Amaral, E.P., Mittereder, L., McCann, K., Correa, A.A.S., Namasivayam, S., Swamydas, M., Moayeri, M., Weiss, J.M., et al. (2022). Mycobacterium tuberculosis Induces Irg1 in Murine Macrophages by a Pathway Involving Both TLR-2 and STING/IFNAR Signaling and Requiring Bacterial Phagocytosis. Front Cell Infect Microbiol 12, 862582 10.3389/fcimb.2022.862582.

53. O’Neill, L.A.J., and Artyomov, M.N. (2019). Itaconate: the poster child of metabolic reprogramming in macrophage function. Nat Rev Immunol 19, 273–281 10.1038/s41577-019-0128-5.

54. Chen, M., Sun, H., Boot, M., Shao, L., Chang, S.J., Wang, W., Lam, T.T., Lara-Tejero, M., Rego, E.H., and Galan, J.E. (2020). Itaconate is an effector of a Rab GTPase cell-autonomous host defense pathway against Salmonella. Science 369, 450–455 10.1126/science.aaz1333.

55. Zhang, B., Lakshmanan, J., Du, Y., Smith, J.W., and Harbrecht, B.G. (2018). Cell-specific regulation of iNOS by AMP-activated protein kinase in primary rat hepatocytes. J Surg Res 221, 104–112 10.1016/j.jss.2017.08.028.

56. Anavi, S., and Tirosh, O. (2020). iNOS as a metabolic enzyme under stress conditions. Free Radic Biol Med 146, 16–35 10.1016/j.freeradbiomed.2019.10.411.

57. Mecsas, J. (2019). Unraveling neutrophil-Yersinia interactions during tissue infection. F1000Res 8 10.12688/f1000research.18940.1.

58. Zhang, Y., Khairallah, C., Sheridan, B.S., van der Velden, A.W.M., and Bliska, J.B. (2018). CCR2(+) Inflammatory Monocytes Are Recruited to Yersinia pseudotuberculosis Pyogranulomas and Dictate Adaptive Responses at the Expense of Innate Immunity during Oral Infection. Infect Immun 86 10.1128/IAI.00782-17.

59. Peterson, L.W., Philip, N.H., DeLaney, A., Wynosky-Dolfi, M.A., Asklof, K., Gray, F., Choa, R., Bjanes, E., Buza, E.L., Hu, B., et al. (2017). RIPK1-dependent apoptosis bypasses pathogen blockade of innate signaling to promote immune defense. J Exp Med 214, 3171–3182 10.1084/jem.20170347.

60. Davis, K.M., Mohammadi, S., and Isberg, R.R. (2015). Community behavior and spatial regulation within a bacterial microcolony in deep tissue sites serves to protect against host attack. Cell Host Microbe 17, 21–31 10.1016/j.chom.2014.11.008.

61. Davis, K.M. (2018). All Yersinia Are Not Created Equal: Phenotypic Adaptation to Distinct Niches Within Mammalian Tissues. Front Cell Infect Microbiol 8, 261 10.3389/fcimb.2018.00261.

62. Lovewell, R.R., Baer, C.E., Mishra, B.B., Smith, C.M., and Sassetti, C.M. (2021). Granulocytes act as a niche for Mycobacterium tuberculosis growth. Mucosal Immunol 14, 229–241 10.1038/s41385-020-0300-z.

63. Huang, L., Nazarova, E.V., Tan, S., Liu, Y., and Russell, D.G. (2018). Growth of Mycobacterium tuberculosis in vivo segregates with host macrophage metabolism and ontogeny. J Exp Med 215, 1135–1152 10.1084/jem.20172020.

64. Dallenga, T., Repnik, U., Corleis, B., Eich, J., Reimer, R., Griffiths, G.W., and Schaible, U.E. (2017). M. tuberculosis-Induced Necrosis of Infected Neutrophils Promotes Bacterial Growth Following Phagocytosis by Macrophages. Cell Host Microbe 22, 519–530 e513 10.1016/j.chom.2017.09.003.

65. Yang, C.T., Cambier, C.J., Davis, J.M., Hall, C.J., Crosier, P.S., and Ramakrishnan, L. (2012). Neutrophils exert protection in the early tuberculous granuloma by oxidative killing of mycobacteria phagocytosed from infected macrophages. Cell Host Microbe 12, 301–312 10.1016/j.chom.2012.07.009.

66. Cohen, S.B., Gern, B.H., Delahaye, J.L., Adams, K.N., Plumlee, C.R., Winkler, J.K., Sherman, D.R., Gerner, M.Y., and Urdahl, K.B. (2018). Alveolar Macrophages Provide an Early Mycobacterium tuberculosis Niche and Initiate Dissemination. Cell Host Microbe 24, 439–446 e434 10.1016/j.chom.2018.08.001.

67. Yaxin Liu, H.Z., Yanjie Hou, Zilin Li, Lin Li, Xiaocui Song, Jingjin Ding, Feng Sha, Yue Xu (2022). Calmodulin Binding Activates Chromobacterium CopC Effector to ADP-Riboxanate Host Apoptotic Caspases. mBio 13 10.1128/mbio.00690-22.

68. Nan Meng, M.X., Ya-Qi Lu, Mi Wang, Krishna M. Boini, Pin-Lan Li and, and Tang, W.-X. (2016). Activation of NLRP3 inflammasomes in mouse hepatic stellate cells during Schistosoma J. infection. Oncotarget 7, 39316–39331 10.18632/oncotarget.10044.

69. Liu, X., Zhang, Y.-R., Cai, C., Ni, X.-Q., Zhu, Q., Ren, J.-L., Chen, Y., Zhang, L.-S., Xue, C.-D., Zhao, J., et al. (2019). Taurine Alleviates Schistosoma-Induced Liver Injury by Inhibiting the TXNIP/NLRP3 Inflammasome Signal Pathway and Pyroptosis. Infection and Immunity 87 10.1128/iai.00732-19.

70. Sanches, R.C.O., Souza, C., Marinho, F.V., Mambelli, F.S., Morais, S.B., Guimaraes, E.S., and Oliveira, S.C. (2020). NLRP6 Plays an Important Role in Early Hepatic Immunopathology Caused by Schistosoma mansoni Infection. Front Immunol 11, 795 10.3389/fimmu.2020.00795.

71. Huppertz, C., Jager, B., Wieczorek, G., Engelhard, P., Oliver, S.J., Bauernfeind, F.G., Littlewood-Evans, A., Welte, T., Hornung, V., and Prasse, A. (2020). The NLRP3 inflammasome pathway is activated in sarcoidosis and involved in granuloma formation. Eur Respir J 55 10.1183/13993003.00119-2019.

72. K Heran Darwin, S.E., José-Carlos Gutierrez-Ramos, Nadine Weich, Carl F Nathan (2003). The proteasome of Mycobacterium tuberculosis is required for resistance to nitric oxide. Science 302, 1963–1966 10.1126/science.1091176.

73. Pichugin, A.V., Yan, B.S., Sloutsky, A., Kobzik, L., and Kramnik, I. (2009). Dominant role of the sst1 locus in pathogenesis of necrotizing lung granulomas during chronic tuberculosis infection and reactivation in genetically resistant hosts. Am J Pathol 174, 2190–2201 10.2353/ajpath.2009.081075.

74. Chacon-Salinas, R., Serafin-Lopez, J., Ramos-Payan, R., Mendez-Aragon, P., Hernandez-Pando, R., Van Soolingen, D., Flores-Romo, L., Estrada-Parra, S., and Estrada-Garcia, I. (2005). Differential pattern of cytokine expression by macrophages infected in vitro with different Mycobacterium tuberculosis genotypes. Clin Exp Immunol 140, 443–449 10.1111/j.1365-2249.2005.02797.x.

75. K Kuida, J.A.L., G Ku, M W Harding, D J Livingston, M S Su, R A Flavell (1995). Altered cytokine export and apoptosis in mice deficient in interleukin-1 beta converting enzyme. Science 267, 2000–2003 10.1126/science.7535475.

76. Rauch, I., Deets, K.A., Ji, D.X., von Moltke, J., Tenthorey, J.L., Lee, A.Y., Philip, N.H., Ayres, J.S., Brodsky, I.E., Gronert, K., et al. (2017). NAIP-NLRC4 Inflammasomes Coordinate Intestinal Epithelial Cell Expulsion with Eicosanoid and IL-18 Release via Activation of Caspase-1 and -8. Immunity 46, 649–659 10.1016/j.immuni.2017.03.016.

77. Dobin, A., Davis, C.A., Schlesinger, F., Drenkow, J., Zaleski, C., Jha, S., Batut, P., Chaisson, M., and Gingeras, T.R. (2013). STAR: ultrafast universal RNA-seq aligner. Bioinformatics 29, 15–21 10.1093/bioinformatics/bts635.

78. Genomics, x. What is Loupe Browser? (Official 10x Genomics Support [Online])https://support.10xgenomics.com/single-cell-gene-expression/software/visualization/latest/what-is-loupe-cell-browser.

79. Satija, R. (2022). Analysis, visualization, and integration of spatial datasets with Seurat https://satijalab.org/seurat/articles/spatial_vignette.html.

80. McInnes, L., Healy, J., and Melville, J. (2020). UMAP: Uniform Manifold Approximation and Projection for Dimension Reduction (arXiv:1802.03426 [stat.ML])doi.org/ 10.48550/arXiv.1802.03426.

81. Tirosh, I., Izar, B., Prakadan, S.M., Wadsworth, M.H., 2nd, Treacy, D., Trombetta, J.J., Rotem, A., Rodman, C., Lian, C., Murphy, G., et al. (2016). Dissecting the multicellular ecosystem of metastatic melanoma by single-cell RNA-seq. Science 352, 189–196 10.1126/science.aad0501.

82. Hao, Y., Hao, S., Andersen-Nissen, E., Mauck, W.M., 3rd, Zheng, S., Butler, A., Lee, M.J., Wilk, A.J., Darby, C., Zager, M., et al. (2021). Integrated analysis of multimodal single-cell data. Cell 184, 3573–3587 e3529 10.1016/j.cell.2021.04.048.

83. Aranda, P.S., LaJoie, D.M., and Jorcyk, C.L. (2012). Bleach gel: a simple agarose gel for analyzing RNA quality. Electrophoresis 33, 366–369 10.1002/elps.201100335.

84. Jian Ye, G.C., Irena Zaretskaya, Ioana Cutcutache, Steve Rozen & Thomas L Madden (2012). Primer-BLAST: A tool to design target-specific primers for polymerase chain reaction. BMC Bioinformatics 13 10.1186/1471-2105-13-134.

85. V E Laubach, E.G.S., O Smithies, P A Sherman (1995). Micelackinginduciblenitricoxidesynthasearenotresistanttolipopolysaccharide-induceddeath. Proc Natl Acad Sci U S A 92, 10688–10692 10.1073/pnas.92.23.10688.

86. Taylor, S.C., Nadeau, K., Abbasi, M., Lachance, C., Nguyen, M., and Fenrich, J. (2019). The Ultimate qPCR Experiment: Producing Publication Quality, Reproducible Data the First Time. Trends Biotechnol 37, 761–774 10.1016/j.tibtech.2018.12.002.

