## Supplementary figures and images for "An innate granuloma eradicates an environmental pathogen using *Gsdmd* and *Nos2*"

### Supplement Figure 1

**Supplement Figure 1.**

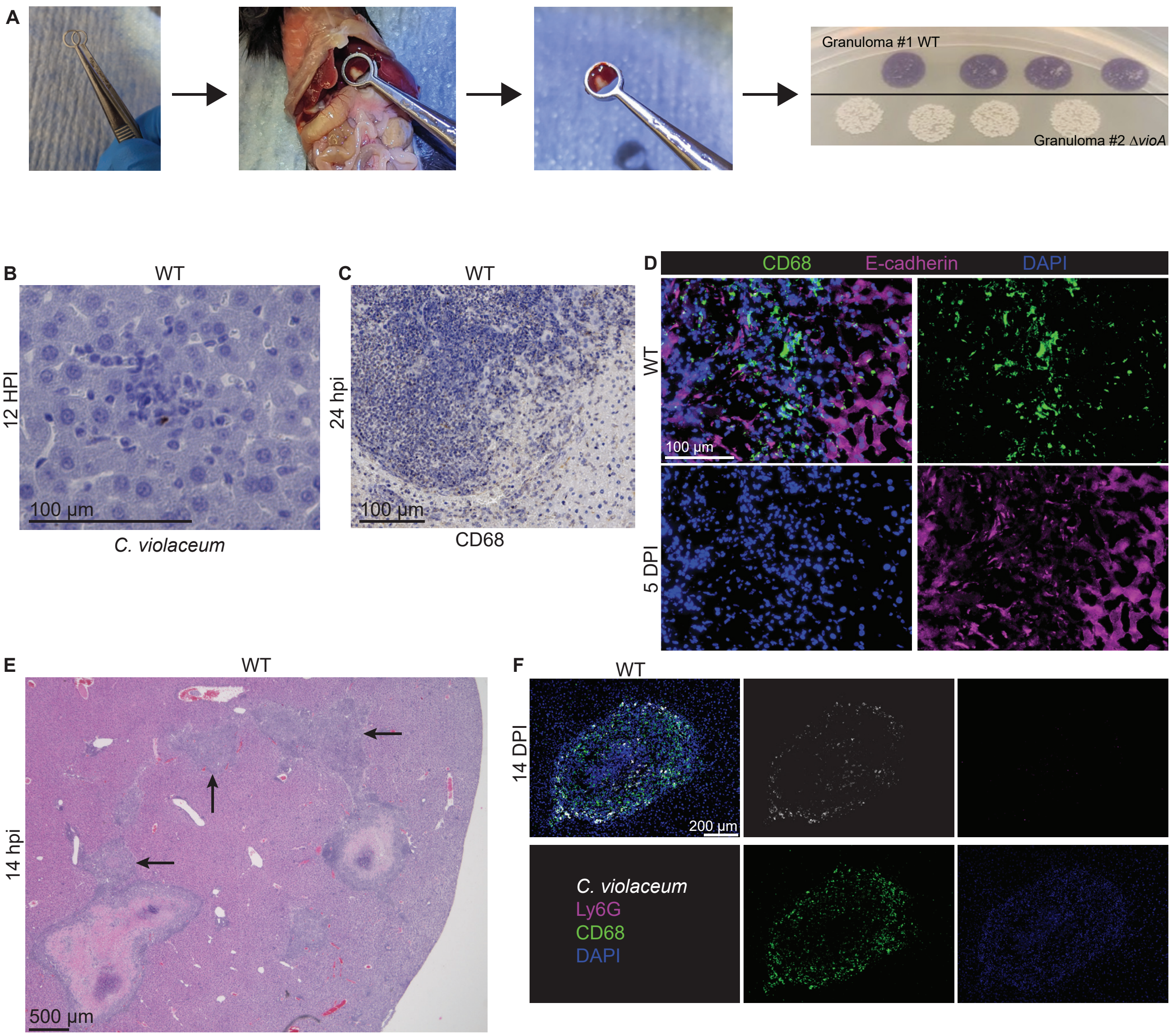

### Supplement Figure 2

##### Supplement Figure 2.

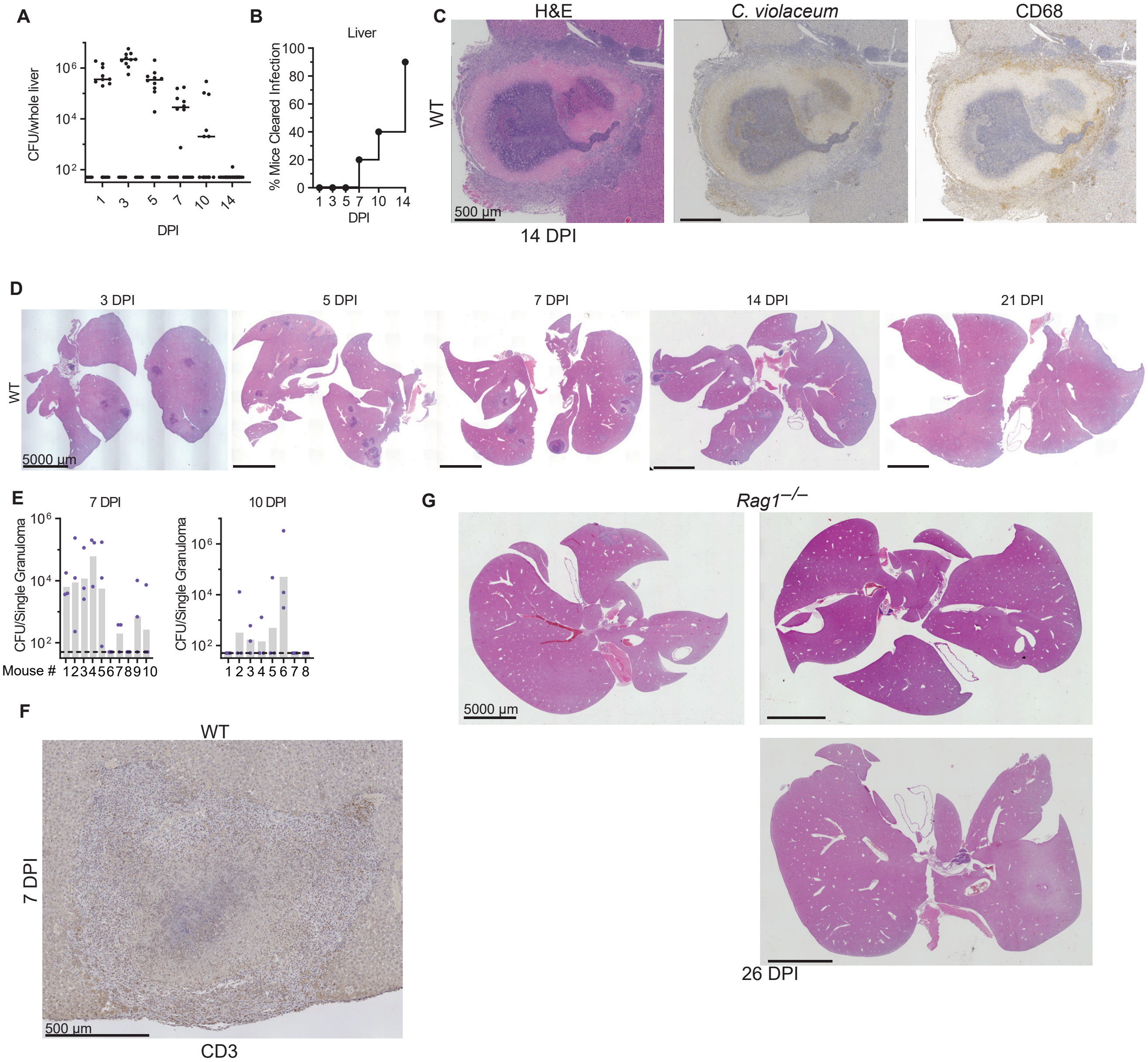

### Supplement Figure 3

**Supplement Figure 3.**

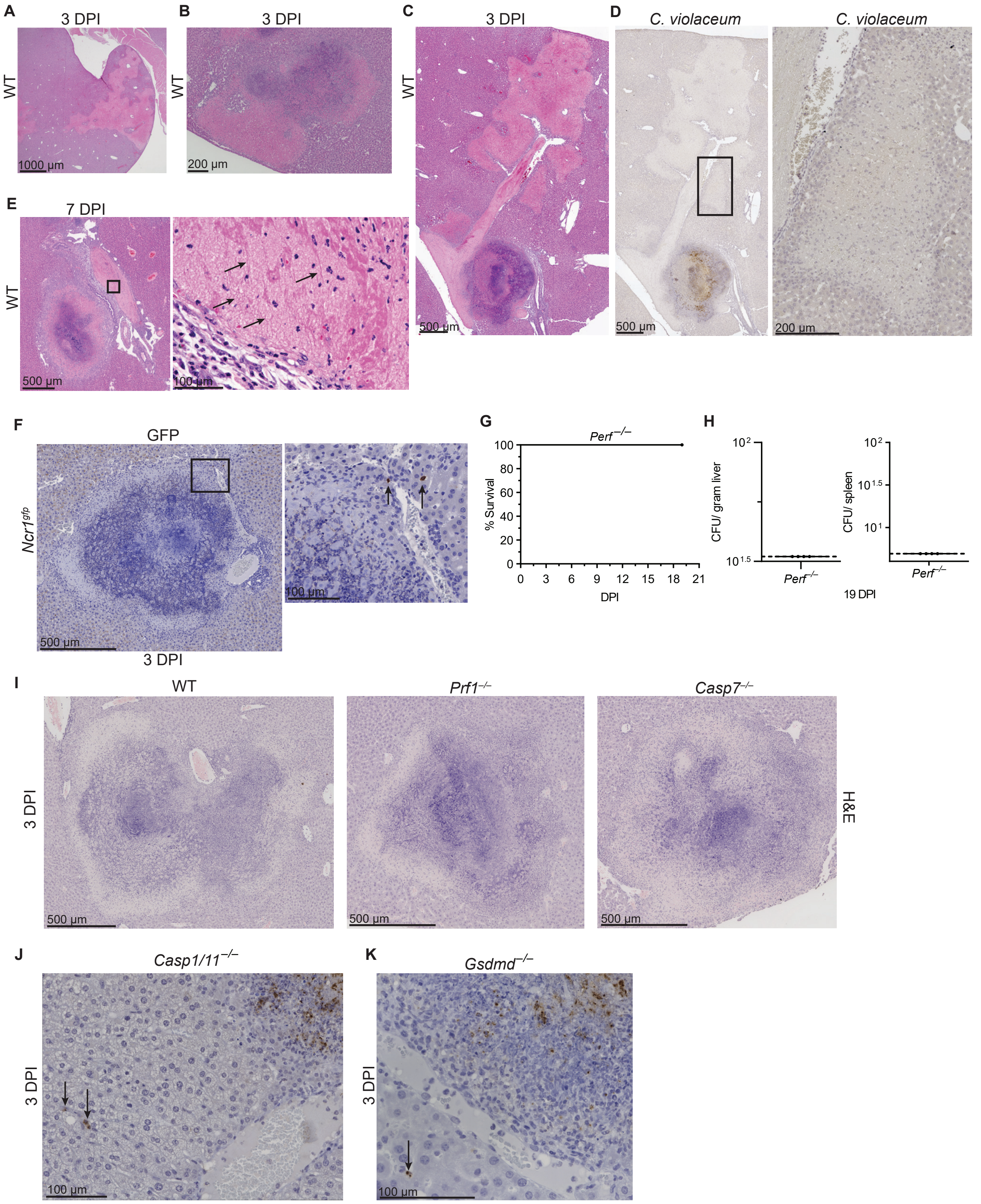

### Supplement Figure 5

#### Supplement Figure 5.

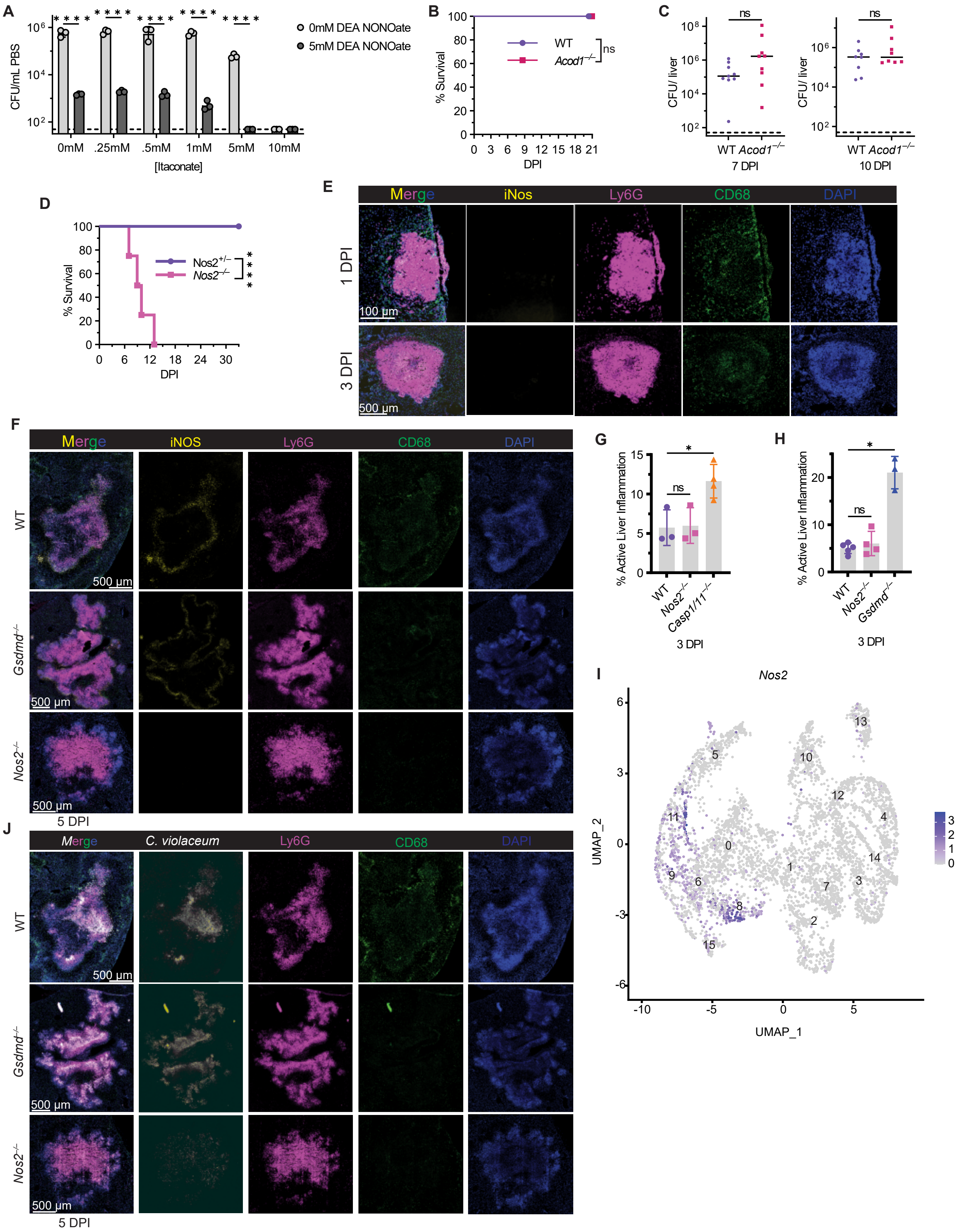
