## Supplement Figure 4 for "An innate granuloma eradicates an environmental pathogen using *Gsdmd* and *Nos2*"

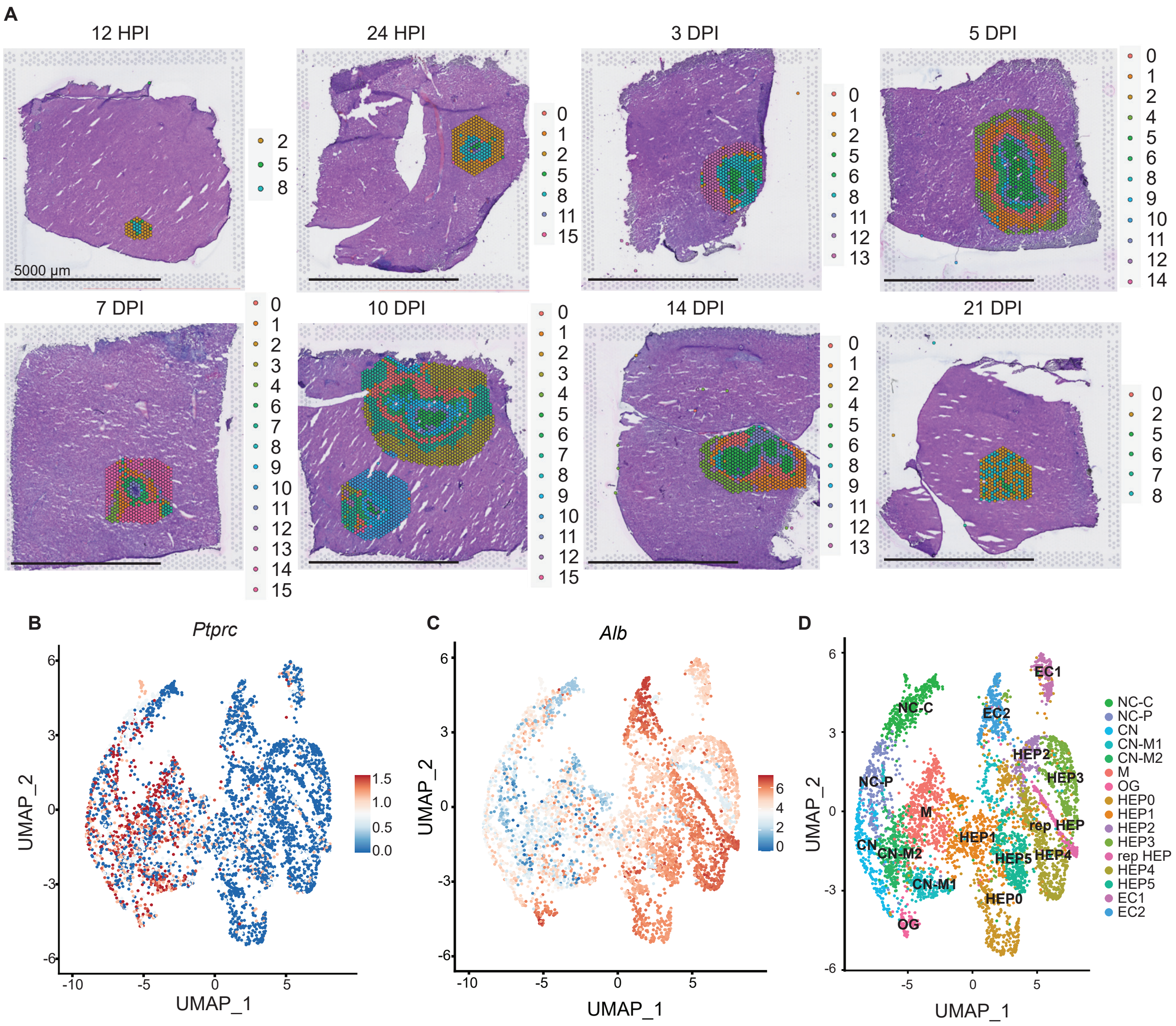


**E**

| Cell Type | Marker Genes Used for Modules |
|---|---|
| Neutrophil | *S100a8, S100a9, Cd33, Csf3r, Ccl3* |
| T Cells | *Cd3d, Cd3e, Cd3g, Trac* |
| Monocytes | *Ly6c2, Ccr2, Fcgr3, Cd14, Itgam* |
| Macrophages | *Aif1, Adgre1, Fcgr1, Cd68, C1qa, C1qb, C1qc* |
| Fibroblasts | *Col1a1, Col1a2, Col3a1, Col5a1, Col5a2* |
| Hepatocytes | *Fabp1, Mup20, Apoa2, Apoc1, Mup3, Apoc3, Scd1, Rbp4, Serpina3k, Alb* |
| Endothelial Cells | *Clec4g, Kdr, Aqp1, Ptprb, Fabp4* |
